# A tumor-secreted protein utilizes glucagon release to cause host wasting

**DOI:** 10.1101/2024.10.24.619567

**Authors:** Guangming Ding, Yingge Li, Chen Cheng, Kai Tan, Yifei Deng, Huiwen Pang, Zhongyuan Wang, Peixuan Dang, Xing Wu, Elisabeth Rushworth, Yufeng Yuan, Zhiyong Yang, Wei Song

**Affiliations:** Department of Hepatobiliary and Pancreatic Surgery, Zhongnan Hospital of Wuhan University, Frontier Science Center for Immunology and Metabolism, Medical Research Institute, Wuhan University, Wuhan, Hubei, China; TaiKang Center for Life and Medical Sciences, Wuhan University, Wuhan, Hubei, China; School of Biomedical Sciences, The University of Queensland, Brisbane, Australia

## Abstract

Tumor-host interaction plays a critical role in malignant tumor-induced organ wasting across multiple species. Despite known regulation of regional wasting of individual peripheral organs by tumors, whether and how tumors utilize critical host catabolic hormone(s) to simultaneously induce systemic host wasting, however, is largely unknown. Using the conserved yki^3SA^-tumor model in *Drosophila*, we discovered that tumors increase the production of adipokinetic hormone (Akh), a glucagon-like catabolic hormone, to cause systemic host wasting, including muscle dysfunction, lipid loss, hyperglycemia, and ovary atrophy. We next integrated RNAi screening and Gal4-LexA dual expression system to identify that yki^3SA^-gut tumors secrete Pvf1 to remotely activate its receptor Pvr in Akh-producing cells (APCs), ultimately promoting Akh production. The underlying molecular mechanisms involved the Pvf1-Pvr axis that triggers Mmp2-dependent ECM remodeling of APCs and enhances innervation from the excitatory cholinergic neurons. Interestingly, we also confirmed the similar mechanisms governing tumor-induced glucagon release and organ wasting in mammals. Blockade of either glucagon or PDGFR (homolog of Pvr) action efficiently ameliorated organ wasting in the presence of malignant tumors. Therefore, our results demonstrate that tumors remotely promote neural-associated Akh/glucagon production via Pvf1-Pvr axis to cause systemic host wasting.

## Introduction

Cancer cachexia, also known as tumor-induced host wasting, is a newly-recognized metabolic disorder that typically involves weight decline, loss of muscle and fat tissues, and hyperglycemia ^1^. Unlike malnutrition, cancer cachexia can hardly be reversed by nutritional supplementation ^2^. Many groups and ours have used rodents, fruit flies, as well as zebrafish, to model tumor-induced host wasting and implicated that, in addition to systemic inflammatory responses ^3^, tumor-secreted factors directly target muscle or adipose tissues and impair energy homeostasis there, leading to muscle wasting or lipid loss ^4–8^. Despite tumor impacts on local metabolism of individual host organs, however, it is not well understood whether and how malignant tumors hijack the essential metabolic hormone(s) from the host to extensively disrupt metabolic homeostasis in multiple organs and lead to systemic host wasting.

We have previously established a cancer-cachexia fly model bearing yki^3SA^ tumors in the gut and further demonstrated that yki^3SA^-gut tumors secrete cachectic ligands, such as Imaginal morphogenesis protein-Late 2 (ImpL2), PDGF- and VEGF-related factor 1 (Pvf1), and Unpaired 3 (Upd3), to lead to energy wasting including lipid loss, muscle dysfunction, ovary atrophy, as well as hyperglycemia, probably via impairment of metabolism and homeostasis of muscle and fat tissues ^9–11^. Importantly, tumor-derived ImpL2/IGFBP2, Upd3/interleukin, and other ligands that result in lipid loss and muscle dysfunction have been consistently found in other tumor-bearing flies and mammals ^12–16^.

*Drosophila* Akh is a well-established homolog of human glucagon that plays conserved roles in mobilization of systemic energy storages ^17,18^. Similar to mammalian glucagon that is produced by pancreatic α-cells to activates glucagon receptor (GcgR) in liver and brain and cause energy depletion ^19,20^, *Drosophila* Akh is produced by neuroendocrine cells in corpora cardiaca (CC) and activates its receptor AkhR in the fat body and certain neurons in the brain to control homeostasis of systemic lipid, carbohydrate and amino acid metabolism. The molecular mechanisms include at least the AkhR-downstream cAMP and Ca^2+^ pathways, which increase glycogenolysis and gluconeogenesis, lipolysis, and amino acid breakdown ^21–25^. Therefore, proper regulation of Akh release is essential for maintenance of organismal energy balance. Previous studies have indicated APCs sense secreted proteins from distal organs (Upd2, NPF, AstA, and AstC) to regulate Akh release and mobilize energy storages under nutrient deprivation ^26–29^. Notably, Akh release is also precisely controlled by upstream inhibitory neurons through release of neurotransmitters or neuropeptides (sNPF and Capa) ^30–32^.

In this study, we characterized that Akh release is required for tumor-induced wasting in *Drosophila* and interestingly uncovered that catabolic ligand Pvf1, which is secreted by yki^3SA^ tumors, directly activates its receptor Pvr in APCs to promotes Akh secretion. We also demonstrated the molecular mechanisms whereby Pvf1-Pvr axis triggers Mmp2-dependent extracellular matrix (ECM) remodeling of APCs and enhances the innervation to upstream excitatory cholinergic neurons. We further validate the conserved regulation of glucagon release in tumor-induced organ wasting in mice.

## Results

### Akh is essential for tumor-induced organ wasting

As Akh is associated with metabolic dysregulation such as lipid loss and hyperglycemia, we wondered whether Akh is involved in systemic organ wasting in yki^3SA^-tumor-bearing flies. To address this hypothesis, we first examined Akh production in the APCs of yki^3SA^-tumor-bearing flies (*esg^TS^>yki^3SA^*). We found that the mass of APCs, intracellular and circulating Akh levels, as well as *Akh* mRNA level, were all increased in yki^3SA^-tumor-bearing flies (**Fig. 1A-E**). We also observed that the expression of *tobi*, an established Akh target gene as indicated by qPCR ^33^, was increased in the yki^3SA^-tumor-bearing flies as compared to control flies (**Fig. 1E**). These data reveal that Akh production is enhanced in yki^3SA^-tumor-bearing flies.

**Figure 1.**
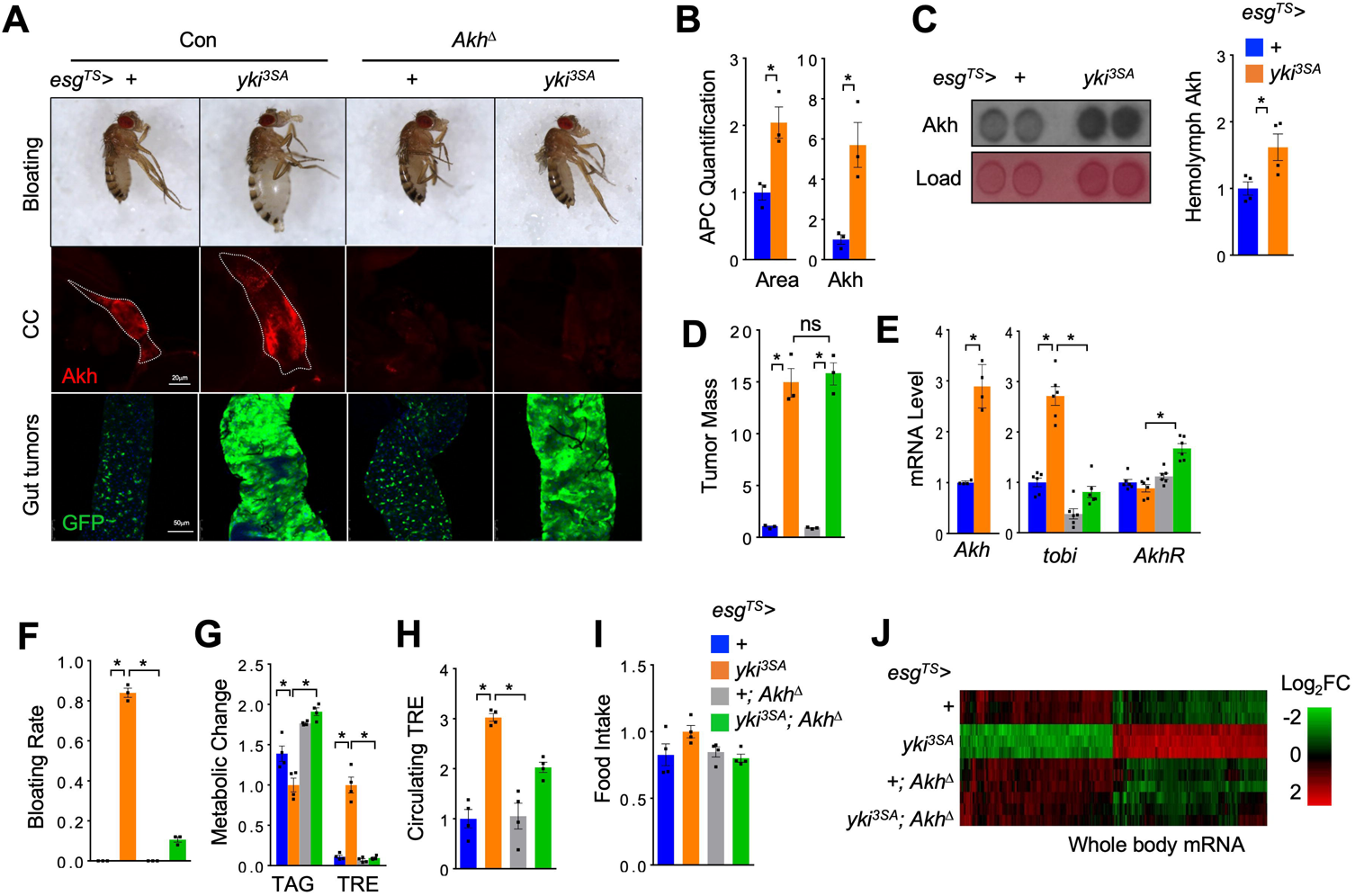
Akh is essential for tumor-induced host wasting in *Drosophila*. Representative images of abdomen bloating (**A,** up), Akh production in APCs (**A,** middle, red, anti-Akh), and gut tumors (**A,** bottom, GFP), quantification of APC masses (area) (**B,** left, n=3) and intracellular Akh amounts (**B,** right, n=3), circulating Akh levels in the hemolymph (**C,** left, dot-blot; right, quantification, n=4), quantification of tumor mass (GFP intensity/gut area) (**D**, n=3), whole body *Akh, AkhR,* and *tobi* mRNA levels (**E,** n=4-6, 5 flies/replicate), bloating rates (**F,** n=3, 20 flies/replicate), metabolic dysregulation such as triglyceride (TAG) and trehalose (TRE) storages (**G**, n=4, 5 flies/replicate), circulating trehalose (TRE) levels in the hemolymph (**H,** n=4, 40 flies/replicate) and food intakes (**I,** n=4, 20 flies/replicate) of adult yki^3SA^-tumor-bearing flies with or without *Akh^Δ^* mutation (*Akh^SAP^/Akh^A^*) at day 8. (**J**) Heatmap indicating both yki^3SA^-tumor- and Akh-dependent differentially expressed genes in the whole-body. Data are presented as mean ± SEM. Each dot represents one biological replicate. Statistical analysis was conducted by two-tailed unpaired t-test (**B, C, E,** left) and one-way ANOVA with Bonferroni’s multiple-comparisons test (**D, E,** right, **F-I**). \**p* < 0.05.

We next generated yki^3SA^-gut tumors in the *Akh*-null mutant flies (*Akh^A/SAP^ or Akh^Δ^*) ^34^, referred as to “*esg^TS^>yki^3SA^*; *Akh^Δ^*”, to examine whether Akh is essential for yki^3SA^-tumor-induced organ wasting. As expected, *Akh* deficiency significantly reduced Akh production and suppressed *tobi* gene expression (**Fig. 1E**). We strikingly observed that *Akh* deficiency robustly improved abdomen bloating, TAG loss, and hyperglycemia (elevation of trehalose, the major circulating carbohydrate composed by two α-glucose in insects) in the yki^3SA^-tumor-bearing flies without affecting the gut-tumor growth or food intake (**Fig. 1A**, **1D-I** and **S1C**). Even though Akh does not directly target muscle or ovary tissues due to absent AkhR expression (single-nucleus RNA-seq data, FlyCellAtlas ^35^), we still found that *Akh* deficiency significantly restored muscle function and ovary homeostasis that were impaired by yki^3SA^ tumors (**Fig. S1A-B**). We next examined the gene expression in the whole body of yki^3SA^-tumor-bearing flies with or without *Akh*-null mutation using RNA-seq and observed that 194 and 201 genes were up- and down-regulated, respectively, by yki^3SA^ tumors in a manner dependent on Akh (**Fig. 1J** and **S1E-F** and **Table. S1**). These differentially regulated genes were found to be enriched in biological processes, pathways, and organelles associated with lipid and carbohydrate metabolism, protein homeostasis, as well as immune responses (**Fig. S1F**). Aligned with amelioration of systemic host wasting by Akh deficiency, these genes are specifically expressed in multiple tissues (**Table. S1**). Notably, the ones expressed in fat body (*Gnmt, CG34136, CG31778, Lsp2*) are implicated in regulation of energy homeostasis and amino acid metabolism, suggesting they are directly targets of Akh signaling (**Table. S1**). Taken together, our results demonstrate that yki^3SA^ tumors remotely promote Akh production to trigger organ wasting.

We also investigated whether *AkhR* deficiency also improved tumor-induced wasting. To do this, we integrated binary expression system ^36,37^ to generate yki^3SA^- gut tumors and knock down *AkhR* expression in the fat body or pan-neurons, two major tissues expressing AkhR (FlyCellAtlas) ^35^, using *LexA* and *GAL4*, respectively. We observed that *AkhR* removal in either tissue at least partially rescued wasting phenotypes including energy loss, abdomen bloating, muscle dysfunction, and ovary atrophy (**Fig. S2**). These results indicate that AkhR in both the fat body and neurons simultaneously contributes to tumor-induced host wasting.

To determine whether Akh gain-of-function in non-tumor control flies sufficiently induce organ wasting, we overexpressed Akh or TrpA1, a heat-activated cation channel that manually promotes peptide hormone release in neurons and endocrinal cells ^29^, in APCs. We found that APC overexpression of TrpA1, but not Akh as reported ^18^, dramatically increased Akh signaling, leading to lipid loss and trehalose elevation (**Fig. S3A-C** and **S3F-H**). However, it failed to cause abdomen bloating, muscle dysfunction or ovary atrophy (**Fig. S3A-J**). Similar outcomes were observed when Akh was ectopically overexpressed Akh in fat body or neurons to manually activate local AkhR signaling in adult flies in an autocrine or paracrine manner (**Fig. S3K-P**) ^18^. These results indicate that excessive Akh release alone is insufficient to cause organ wasting, despite its roles in carbo-lipid metabolic mobilization.

### The receptor Pvr promotes Akh release in APCs

We hypothesized that yki^3SA^ tumors secrete cachectic protein(s) to target specific receptors in APCs and regulate Akh release (**Fig. 2A**). To identify these regulatory receptors, we performed an *in vivo* RNAi screening against transmembrane proteins in the larval APCs by crossing 311 RNAi lines to *Akh-GAL4* and measured glycemic changes (circulating trehalose) (**Table. S2**), a quick and reliable readout of larval Akh response ^21^. As expected, RNAi against either *Akh* or *AstA-R2*, which was found to promote Akh production ^28^, significantly suppressed glycemic level as compared to control RNAi (*w-i)* (**Fig. 2B**). We eventually found 22 and 38 RNAi lines in APCs that down- and up-regulated glycemic levels by >10%, respectively (**Fig. S4A**). These hits included receptors for brain-gut hormones, neurotransmitters, olfactory and gustatory molecules, as well as ion and mechanical channels, suggesting a comprehensive network in APCs that modulates Akh release and glycemic homeostasis (**Fig. S4A** and **Table. S2**). Interestingly, RNAi against *PDGF- and VEGF-receptor related* (*Pvr*), an RTK specifically activated by yki^3SA^ tumor-derived Pvf1 ^38^, was a top hit in APCs that dramatically decreased glycemic level (circulating trehalose) (**Fig. 2B**). To validate effects of Pvr in APCs, we examined the expression pattern of Pvr using an endogenous GAL4 line and confirmed *Pvr-GAL4*-driven GFP expression in larval APCs (**Fig. S4B**). In addition to *Pvr* RNAi, overexpression of a dominant negative form of Pvr (*Pvr^DN^*) in APCs also suppressed larval glycemic level (**Fig. S4C**). These results indicate that Pvr in the larval APCs promotes Akh release.

**Figure 2.**
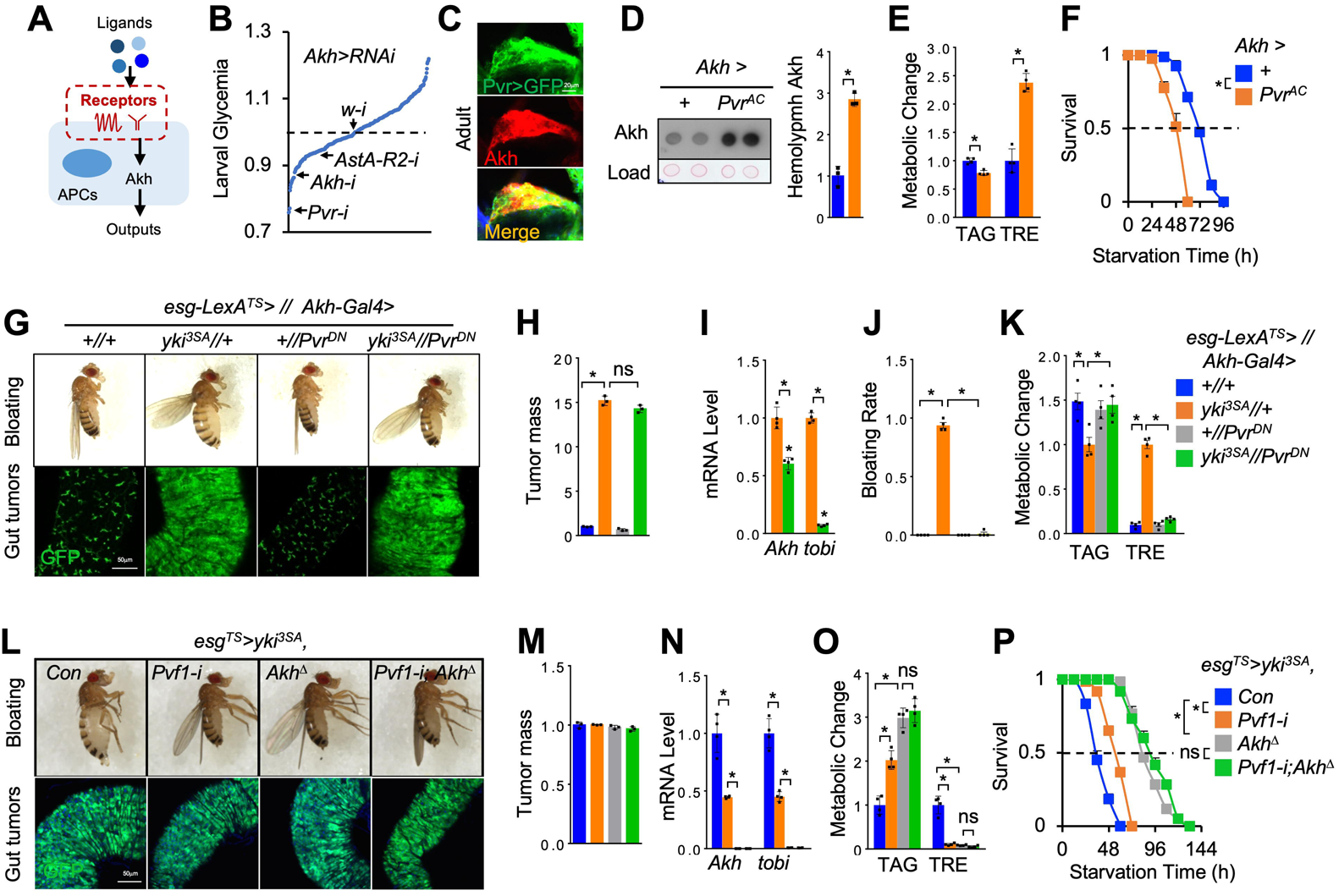
Pvr signaling regulates Akh release. (**A-B**) Experimental strategy (**A**) and glycemic changes (**B**) of the *in vivo* RNAi screening against transmembrane proteins in the larval APCs. (**C**) Immunostaining indicating Pvr expression (green, Pvr>GFP; red, anti-Akh) in adult APCs. (**D-F**) Circulating Akh levels in the hemolymph (**D,** left, dot-blot; right, quantification, n=3), metabolic changes such as TAG and trehalose (TRE) storages (**E**, n=3, 5 flies/replicate), as well as survival rates under starvation (**F**, n=4, 20 flies/replicate), of adult flies with Pvr^AC^ overexpression in APCs at day 4. (**G-N**) Bloating phenotype (up) and gut-tumors (bottom, green) (**G, L**), quantification of tumor mass (GFP intensity/gut area) (**H, M,** n=3), whole body *Akh* and *tobi* expression (**I**, **N,** n=3, 5 flies/replicate), wasting effects such as bloating rates (**J,** n=3, 20 flies/replicate), TAG and TRE storages (**K**, **O**, n=3, 5 flies/replicate), and starvation resistance (**P**, n=4, 20 flies/replicate) of yki^3SA^- tumor-bearing flies with genetic manipulation in APCs at day 6 (**G-K**, *LexA+GAL4*) or with tumor *Pvf1* knockdown or systemic *Akh* deficiency (*Akh^Δ^*, *Akh^SAP^/Akh^A^*) at day 8 (**L-P**, *GAL4*). Data are presented as mean ± SEM. Each dot represents one biological replicate. Statistical analysis was conducted by two-tailed unpaired t-test (**D, E**), one-way ANOVA with Bonferroni’s multiple-comparisons test (**H-K, L-O**), or log-rank test (**F, P**). \**p* < 0.05.

We next studied the Pvr functions in adult flies. The *Pvr-GAL4* line also confirmed the Pvr expression in adult APCs (**Fig. 2C**). We also found that overexpression of a constitutively active Pvr (*Pvr^AC^*) in adult APCs increased Akh production, including circulating Akh level in the hemolymph and systemic mRNA levels of both *Akh* and *tobi*, enhanced Ca^2+^ signaling in APCs, and perturbed carbo-lipid metabolism, including whole-body TAG and trehalose level and survival under starvation (**Fig. 2D-F, S1D, S4C, S4D** and **S4F**). In contrast, flies with *Pvr^DN^* overexpression or *Pvr* RNAi in APCs exhibited opposite phenotypes (**Fig. S4E-H**). To avoid developmental effects, we further used *tub-Gal80^TS^* to manipulate Pvr in adult APCs only. Consistently, APC Pvr activation for 4 days sufficiently increased Akh production and energy catabolism, while adult Pvr inactivation showed relatively mild effects (**Fig. S4I-L**). Finally, *Akh* knockdown in the context of Pvr^AC^ overexpression in the APCs dramatically alleviates energy catabolism and restored Akh response (**Fig. S7A-C**), indicating an Akh-dependent metabolic role of Pvr. Taken together, these results demonstrate that Pvr signaling in APCs promotes Akh production.

### Tumor-derived Pvf1 activate Pvr in APCs to enhance Akh release

To verify the physiological effects of Pvr-associated Akh release in the context of yki^3SA^-gut tumors, we expressed Pvr^DN^ using *Akh*-*GAL4* to inactivate Pvr specifically in APCs of yki^3SA^-tumor-bearing flies. Strikingly, as compared to marginal effects of APC Pvr inactivation in non-tumor adult flies, we observed that Pvr inactivation in APCs of yki^3SA^-tumor-bearing flies resulted in a significant decrease in Akh response (*Akh* and *tobi* expression) and robustly improved host wasting, including bloating, lipid loss, hyperglycemia, muscle dysfunction, and ovary atrophy without affecting tumor growth at day 6 (**Fig. 2G-K** and **S5A-B**). These results were further confirmed by knocking down *Pvr* expression in APCs of yki^3SA^-tumor-bearing flies (**Fig. S5C-G**).

Given the distribution Pvr in muscle and adipose tissue ^11^, we wondered whether adipose or muscle Pvr signaling also contributes to host wasting. We expressed Pvr^DN^ in muscle and fat body using *Mhc-* and *R4-GAL4*, respectively, in the yki^3SA^- tumor-bearing flies. However, Pvr inactivation in neither fat body nor muscle improved yki^3SA^-tumor-induced host wasting (**Fig. S6**), except for a slight rescue in abdomen bloating and hyperglycemia by fat body Pvr inactivation (**Fig. S6C** and **S6E**). Thus, our data demonstrate that yki^3SA^ tumors cause host wasting predominantly through Pvr function in APCs.

To examine whether tumor-derived Pvf1 functions through Akh production, we performed the genetic interaction between *Pvf1* and *Akh* in yki^3SA^-tumor-bearing flies. We found that either *Akh*-null mutation or tumor-specific *Pvf1* knockdown potently diminished wasting effects, including bloating, lipid loss, hyperglycemia, as well as starvation sensitivity, in yki^3SA^-tumor-bearing without affecting tumor growth at day 8 (**Fig. 2L-P**). Tumor-specific *Pvf1* knockdown also significantly decreased both *Akh* and *tobi* mRNA levels, albeit to a less extent than *Akh*-null mutation (**Fig. 2N**). Importantly, *Akh* mutation plus tumor-*Pvf1* knockdown failed to further significantly alleviate wasting effects as compared to single one(s) (**Fig. 2L-P**). These results demonstrate that tumor-derived Pvf1 functions, at least partially, through Akh to cause host wasting.

### Pvf1/Pvr axis promotes Akh release via ERK/Mmp2 signaling

Previous studies showed that Pvr activates Ras/Raf/MEK/ERK and multiple downstream targets to regulate various events of tissue homeostasis, such as cell proliferation, differentiation, as well as migration ^38^. To investigate the molecular mechanisms of Pvr regulation of Akh release, we overexpressed Pvr^AC^ to activate Pvr signaling in APCs and simultaneously knocked down potential downstream targets. We interestingly observed that, in the context of APC Pvr^AC^ overexpression, *ERK* knockdown significantly increased fly survival rates under starvation, suppressed both *Akh* and *tobi* gene expression, increased TAG levels, and decreased trehalose levels (**Fig. 3A-B** and **S7D**). On the other hand, overexpression of an active Raf (*Raf^F179^*) to activate MEK/ERK signaling in the APCs of wild-type flies phenocopied the effects of Pvr activation to promote Akh release, enhance systemic Akh response, and impaired carbo-lipid metabolic homeostasis (**Fig. S7E- G**). We further performed RNAi screening of ERK downstream targets and, strikingly, found that knockdown of *Matrix metalloproteinase 2* (*Mmp2*), but not *Mmp1* ^39,40^, potently rescued Pvr^AC^-associated starvation sensitivity (**Fig. 3A**). Biochemical and metabolic analysis consistently revealed that *Mmp2* deficiency suppressed Akh release, decreased *tobi* expression, as well as abolished lipid loss and hyperglycemia (**Fig. 3B-C**, **S7D** and **S7H-I**). Overexpression of Timp, a single homolog of the tissue inhibitors of metalloproteinases (TIMPs) blocking Mmp1/2 in fly ^41^, in the context of Pvr^AC^ overexpression in APCs also suppressed Akh release and alleviated subsequent signaling and metabolic outputs (**Fig. 3A-C** and **S7D**). Note that, neither *Mmp2* knockdown or *Timp* overexpression in APCs of control flies affected systemic Akh signaling or carbo-lipid metabolism (**Fig. S7J-L**), demonstrating their roles in regulating Akh release are contingent upon Pvr activation. APC ERK knockdown in control flies decreased Akh signaling and carbo-lipid mobilization (**Fig. S7J-L**), suggesting that ERK might function through Mmp2 and other targets.

**Figure 3.**
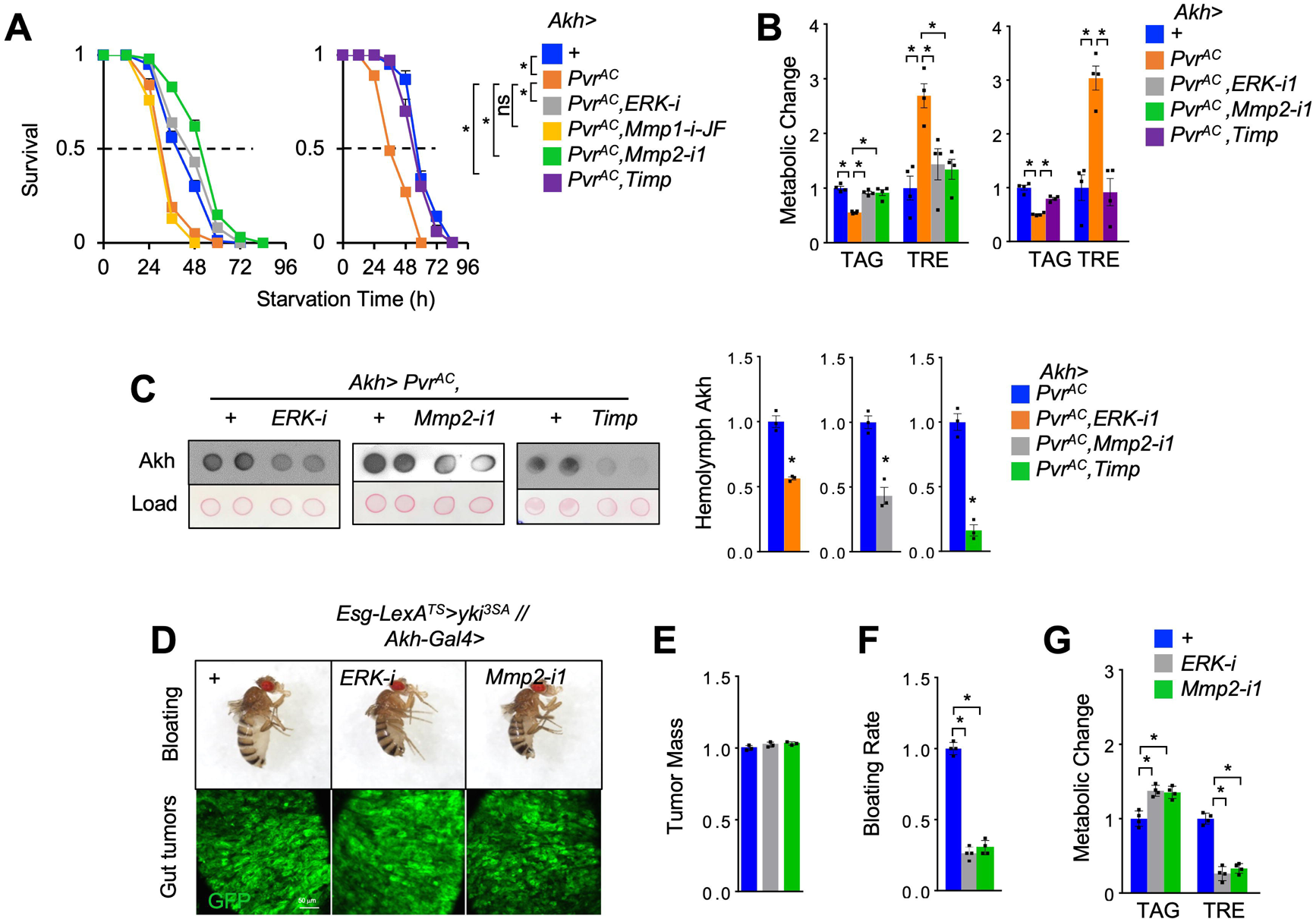
Pvr enhances Akh release via ERK/Mmp2 signaling. (**A-C**) Survival rates under starvation (**A**, n=4, 20 flies/replicate), metabolic changes such as TAG and trehalose (TRE) storages (**B**, n=4, 5 flies/replicate), and hemolymph Akh levels (**C,** left, dot-blot; right, quantification, n=3) of adult flies bearing indicated RNAi in the context of Pvr^AC^ overexpression in APCs at day 4. (**D-G**) Bloating phenotype (up) and gut-tumors (bottom, green) (**D**), quantification of tumor mass (GFP intensity/gut area) (**E,** n=3), bloating rates (**F,** n=4, 20 flies/replicate), and TAG and TRE storages (**G**, n=4, 5 flies/replicate) of yki^3SA^-tumor-bearing flies with genetic manipulation in APCs at day 6 (*LexA+GAL4*). Data are presented as mean ± SEM. Each dot represents one biological replicate. Statistical analysis was conducted by two-tailed unpaired t-test (**C**), one-way ANOVA with Bonferroni’s multiple-comparisons test (**B, E-G**), or log-rank test (**A**). \**p* < 0.05.

We next validated the wasting effects of ERK/Mmp2 axis in APCs in the context of yki^3SA^-gut tumors. Using *GAL4/LexA* binary system, we knocked down expression of *ERK* or *Mmp2* in APCs of yki^3SA^-tumor-bearing flies and consistently found that host wasting, including bloating, lipid loss, and hyperglycemia, were robustly improved without affecting yki^3SA^ tumors in the gut at day 6 (**Fig. 3D-G**). Taken together, these results demonstrate that Pvr-ERK-Mmp2 axis enhances Akh release to cause energy wasting in yki^3SA^-tumor flies.

### Pvf1-Pvr axis enhances Mmp2-dependent ECM remodeling and neuronal innervation of APCs

We next investigate the molecular mechanisms by which Mmp2 regulates Akh production. Mmp-mediated homeostasis of ECM is essential for multiple neuronal activities, including neuronal plasticity, synaptic formation and neural innervation ^42^. APCs have been reported to be targeted by upstream regulatory neurons to control Akh release ^30,31^. We therefore wonder whether Pvr regulates Akh release via Mmp2-induced ECM remodeling and neural contact. To address it, we first accessed the ECM homeostasis in the APCs using GFP-tagged collagen IV (Viking, Vkg-GFP) and to label ECM ^43^. Interestingly, we observed very strong Vkg-GFP signals around somas of >20 APCs in wild-type adult flies (**Fig. 4A**). Pvr^AC^ overexpression in APCs potently decreased, whereas *Mmp2* RNAi in the context of Pvr^AC^ restored, Vkg-GFP signals around APCs at day 4 (**Fig. 4A** and **4C**), indicating Pvr-Mmp2 regulation of ECM remodeling. Consistent with Pvr activation by tumor-derived Pvf1, similar patterns of ECM in APCs were observed in flies bearing yki^3SA^ tumors with or without Pvf1 knockdown at day 8 (**Fig. 4B** and **4D**). Similar results were observed using another ECM indicator, integrin βPS (integrin) (**Fig. S8A-B**) ^43^.

**Figure 4.**
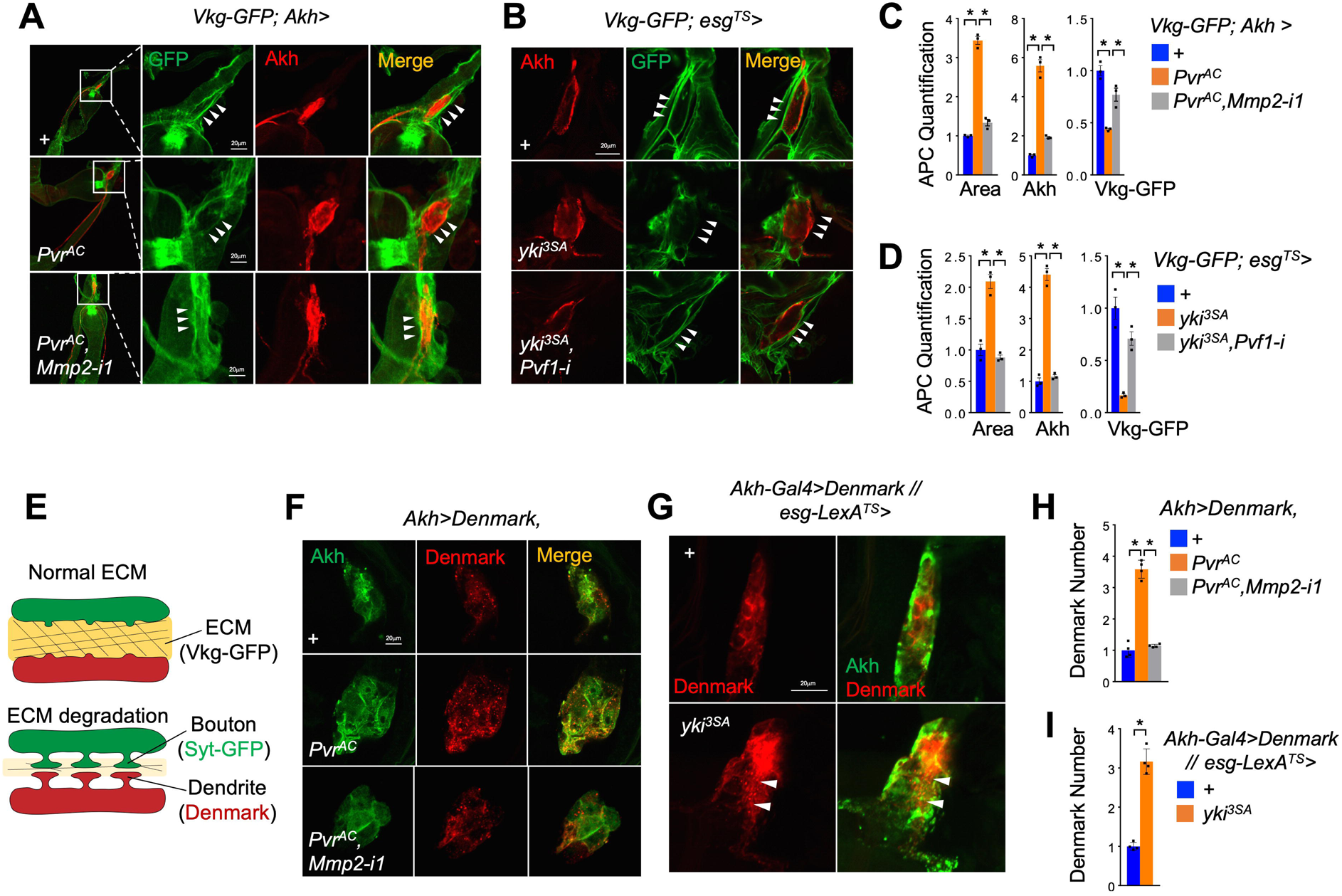
Pvf1/Pvr axis enhances neural contacts of APCs via ECM remodeling. (**A-D**) ECM homeostasis indicated by Vkg-GFP around the somas of APCs (**A**, **B,** green, GFP; red, anti-Akh) and quantification of APC masses (area), intracellular Akh amounts, as well as extracellular Vkg-GFP amounts (**C, D,** n=3) of adult flies with Pvr^AC^ overexpression plus *Mmp2* RNAi in APCs at day 4 or flies bearing yki^3SA^-tumor plus *Pvf1 R*NAi at day 8. (**E**) The model showing that ECM (Vkg-GFP) degradation promotes synapse contact including bouton (Syt-GFP) and dendrite (Denmark) formation. (**F-J**) Dendrites labeled by Denmark (**F, G**, red; green, anti-Akh) and quantification of dendrite numbers (**H, I,** n=4) in APCs of adult flies with Pvr^AC^ overexpression plus *Mmp2* RNAi in APCs at day 4 or yki^3SA-^tumor-bearing flies (*LexA+GAL4*) at day 6. Data are presented as mean ± SEM. Each dot represents one biological replicate. Statistical analysis was conducted by two-tailed unpaired t-test (**I**) or one-way ANOVA with Bonferroni’s multiple-comparisons test (**C, D, H**). \**p* < 0.05.

As ECM remodeling promotes synapse formation (**Fig. 4E**), we next examined the synaptic contact using Dendritic Marker (Denmark) to visualize dendrites in the APCs ^44^. APC Pvr^AC^ overexpression consistently increased dendrite numbers as indicated by Denmark inside APCs, APC masses, and Akh production in an Mmp2-dependent manner at day 4 (**Fig. 4F** and **4H**). yki^3SA^-tumor-bearing flies also exhibited an increase in dendrite numbers of APCs as indicated by Denmark puncta (**Fig. 4G** and **4I**). These results indicate that Pvf1-Pvr-Mmp2 axis modulates ECM and increases neural innervation of APCs in yki^3SA^-tumor-bearing flies.

### Pvf1-Pvr axis promotes cholinergic innervation of APCs to increase Akh release

We next exploit the potential excitatory innervating neuron(s) upstream of APCs by screening different neurotransmitter-producing neurons, including cholinergic (*Cha-GAL4*), dopaminergic (*TH-GAL4*), 5-HT (*TRH-Gal4*), octopaminergic (*Tdc2-GAL4*), GABAergic (*Gad1-GAL4*), and glutamatergic (*VGlut-GAL4*) neurons ^45^. We expressed a presynaptic marker, GFP-tagged Synaptotagmin (Syt-GFP), to visualize boutons in these neurons and found strong Syt-GFP signals driven by, at least, *Cha-GAL4* in the APCs (**Fig. 5A** and **S8C**). Meanwhile, we found that APCs express multiple receptors for acetylcholine from published single-nucleus RNA-seq dataset (**Fig. S8D**) ^46^. This cholinergic-APC innervation was further confirmed by the trans-tango system (**Fig. 5B**). We next thermally activated cholinergic neurons by overexpressing *TrpA1* at 29 °C and found enhanced systemic Akh response as indicated by *tobi* expression, as well as lipid loss and hyperglycemia, in the adult flies (**Fig. 5C**). These data indicate that, at least, cholinergic neurons functionally projection onto APCs.

**Figure 5.**
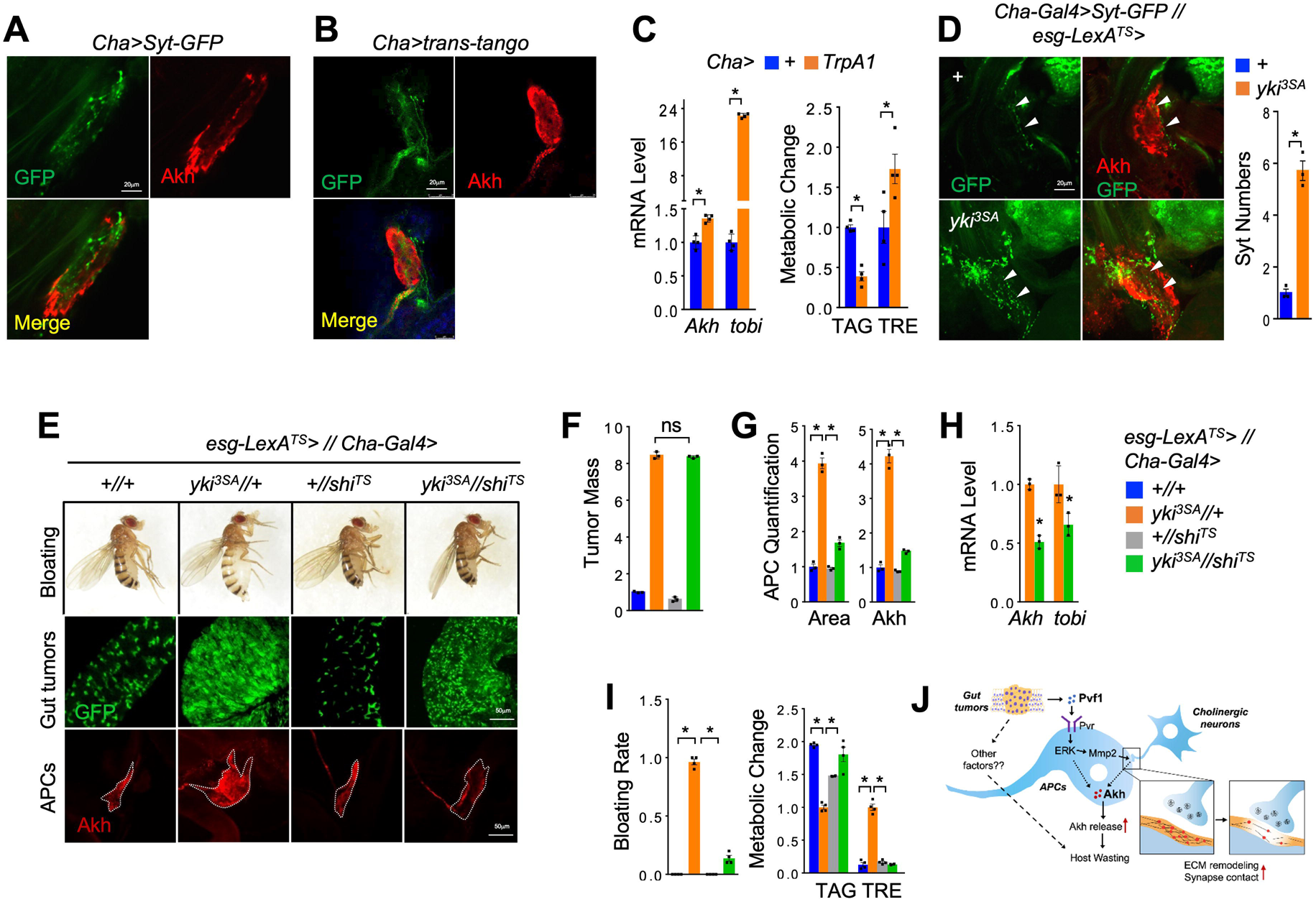
Cholinergic innervation of APCs is essential for yki^3SA^-tumor-induced Akh release and host wasting. (**A**) Cholinergic boutons indicated by Cha>Syt-GFP (green) in the somas of adult APCs (red, anti-Akh) at day 4. (**B**) Trans-tango system indicating overlaps between cholinergic innervating neurons (GFP) and APCs (red, anti-Akh). (**C**) Whole body *Akh* and *tobi* expression (left, n=3, 5 flies/replicate) and metabolic changes such as TAG and trehalose (TRE) storages (right, n=4, 5 flies/replicate) of adult flies with activation of cholinergic neurons induced at 29°C for 12 hours. (**D**) Upstream cholinergic boutons indicated by Syt-GFP (left, green) in APCs (left, red, anti-Akh) and quantification of bouton numbers (right, n=3) of yki^3SA^-tumor-bearing flies (*LexA+GAL4*) at day 6. (**E-H**) Bloating phenotype (**E,** up), gut tumors (**E,** middle, green) and quantification (**F,** n=3), and APCs (**E,** bottom, red, anti-Akh), quantification of APC masses (area) and intracellular Akh amounts (**G**, n=3), whole body *Akh* and *tobi* expression (**H**, n=3, 5 flies/replicate), abdomen bloating rates (**I,** left, n=4) and TAG and TRE storages (**I,** right, n=4, 5 flies/replicate) of yki^3SA^-tumor-bearing flies with synapse disruption in cholinergic neurons at day 6 (*LexA+GAL4*). (**J**) The schematic model illustrating the release of Akh and its regulation by tumor-secreted Pvf1 and ECM-associated neural innervation in yki^3SA^-tumor-bearing flies, leading to organ wasting. Data are presented as mean ± SEM. Each dot represents one biological replicate. Statistical analysis was conducted by two-tailed unpaired t-test (**C, D**), one-way ANOVA with Bonferroni’s multiple-comparisons test (**F- I**). \**p* < 0.05.

We further observed a significant increase in the number of Syt-GFP-labeled cholinergic boutons in the APCs of flies bearing yki^3SA^ tumors at day 8 (**Fig. 5D**). Finally, to evaluate the functional impacts of cholinergic projections on APCs, we thermally disrupted neurotransmission of cholinergic neurons by expressing a temperature-sensitive mutant of *shibire* (*shi^TS^*), the fly homolog of dynamin essential for synaptic vesicle recycling in nerve terminals ^47^, at 29 °C. Interestingly, *shi^TS^* expression in cholinergic neurons of yki^3SA^-tumor-bearing flies significantly suppressed Akh production and alleviated host wasting like bloating, lipid loss, and hyperglycemia at day 6 (**Fig. 5E-I**). Taken together, our data demonstrate that cholinergic innervation of APCs is increased to cause energy wasting in yki^3SA^-tumor-bearing flies.

### Glucagon is required for tumor-induced wasting in mammals

Since fly Akh and mammalian glucagon are conserved catabolic hormones, we wonder whether neural-associated glucagon release also participates in tumor-induced host wasting in mammals. To do this, we first assessed the glucagon changes in tumor-bearing mammals. We measured the circulating glucagon levels in the patients bearing pancreatic cancer that is with > 85% chance to develop carbo-lipid wasting and weight decline ^48^. Glucagonoma a rare tumor that produces high levels of glucagon, was excluded in advance in this study. Interestingly, as compared to pancreatic benign diseases (n=15), patients bearing pancreatic cancer without (n=18) or with weight decline (>1.5% per month) (n=21) within three months prior to surgery potently exhibited higher serum glucagon concentration (**Fig. 6A** and **Table. S3**). Considering the high prevalence of weight loss and body-composition change among pancreatic cancer patients in the end, it is possible that hyperglucagonemia might occur prior to the onset of weight loss or organ wasting. We also observed increased mass of α-cells in normal pancreatic tissues of cancer patients (**Fig. 6B**).

**Figure 6.**
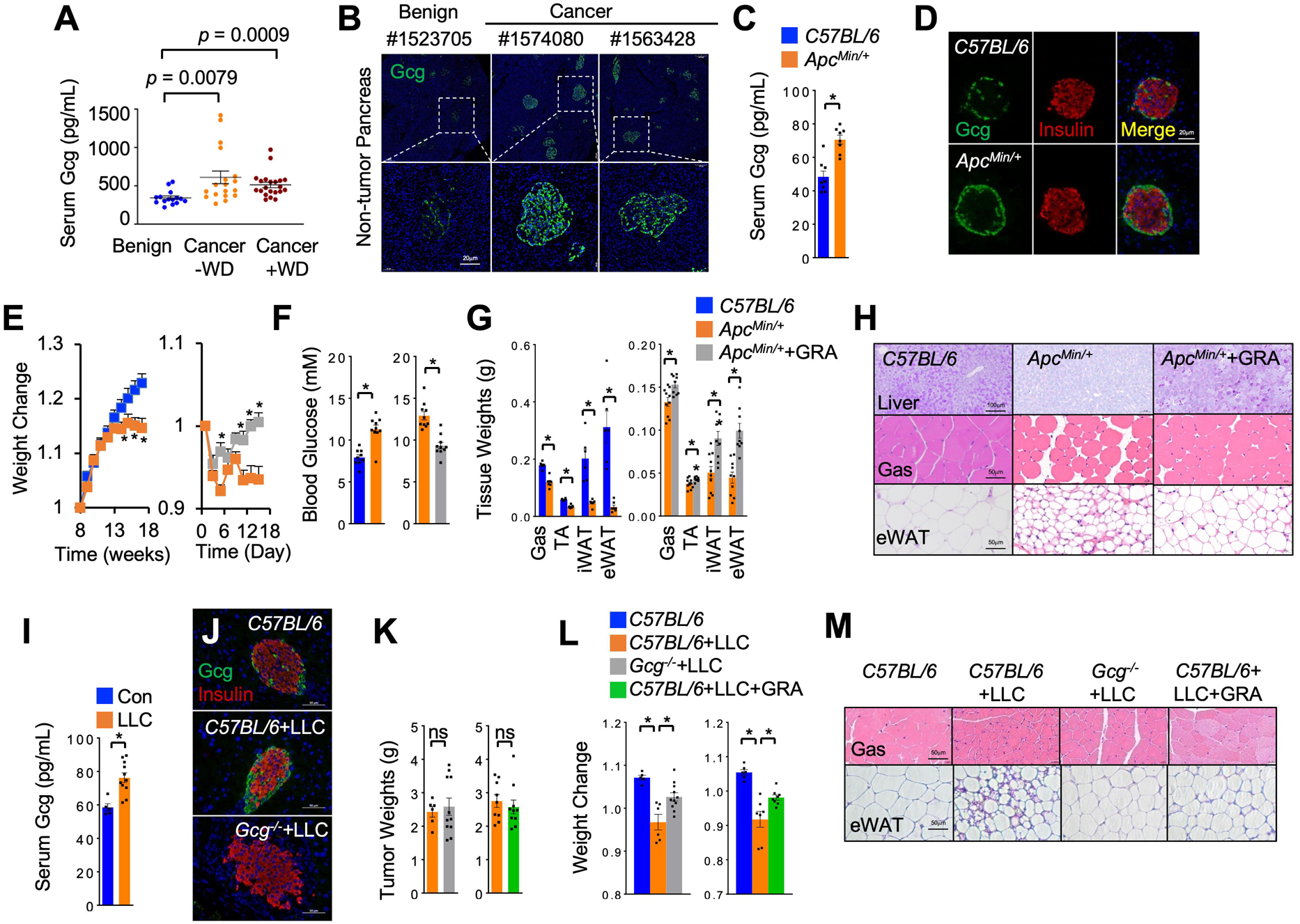
Excessive glucagon release is essential for tumor-induced wasting in mammals. (**A**) Circulating glucagon levels in patients with benign pancreatic disease (n=15) and pancreatic-cancer patients with (n=21) or without (n=18) weight decline. (**B**) Representative islet morphologies (green, Gcg) indicated by confocal images in non-tumor tissue of patients. (**C-D**) Serum glucagon levels (**C**) and islet morphologies indicated by confocal images (**D**, green, anti-Gcg; red, anti-insulin) of *Apc^Min/+^* mice after 18 weeks (n=8). (**E-H**) Whole body weight changes (**E**), fed blood glucose (**F**), tissue weights (**G**), liver glycogen contents (**H**, PAS staining) and tissue morphologies (**H**, Gas, myotube diameters; eWAT, adipocyte sizes) of *Apc^Min/+^* mice from week 18 with or without daily IP injection of 10 mg/Kg/day GRA Ex-25 for two weeks (n=10). (**I-M**) Serum glucagon levels (**I**) islet morphologies (green, anti-glucagon; red, anti-insulin) (**J**), tumor weights (**K**), body weight changes (**L**), and tissue morphologies (**M**, Gas, myotube diameters; eWAT, adipocyte sizes) of indicated LLC-tumor-bearing mice (*C57BL/6*, n=5; *C57BL/6*+LLC, n=6; *Gcg^-/-^+*LLC, n=10) or LLC-tumor-bearing mice with daily IP injection of inhibitors from day 14 for 7 days (PBS, n=6; LLC, n=7; LLC+GRA, n=8, 10 mg/Kg/day). Data are presented as mean ± SEM. Each dot represents one biological replicate. Statistical analysis was conducted by two-tailed unpaired t-test (**C, E-G, I, K**), one-way ANOVA with Bonferroni’s multiple-comparisons test (**A, L**). \**p* < 0.05.

We next examined the glucagon levels in tumor-bearing mice. *Apc^Min/+^* mice are an established colon-cancer model that exhibits weight decline, fat loss, and muscle atrophy at 16-18 weeks ^49,50^. In line with human observations, we found elevated serum glucagon levels and enlarged α-cells in *Apc^Min/+^* mice when they started showing wasting symptoms (**Fig. 6C-E**). We analyzed the carbo-lipid metabolism in cachectic *Apc^Min/+^* mice after 18 weeks and observed upregulation of glucagon-target genes in the liver, glucose intolerance, higher blood glucose levels, as well as hepatic glycogen depletion (**Fig. 6F-H** and **S9A-B**), as compared to age-matched wild-type *C57BL/6* mice.

Previous results have indicated that excessive glucagon response potently also results in weight decline, lipid loss, and muscle atrophy in mice ^51–55^, we next investigate whether glucagon is essential for tumor-induced systemic wasting. We generated *Gcg*-null mice with removal of exon 3-6 that contains glucagon coding region and crossed them to *Apc^Min/+^* lines to obtain *Apc^Min/+^*; *Gcg^-/-^* mice. However, *Apc^Min/+^*; *Gcg^-/-^*mice appeared very unhealthy and started dying within week 15 with unknown mechanism (data not shown). We thus intraperitoneally (IP) injected small-molecule inhibitors, GRA Ex25 (GRA), of GCGR in *Apc^Min/+^* mice after 18 weeks to suppress glucagon response instead. Interestingly, injection of GRA for 7 days significantly alleviated weight loss, glucose intolerance, and hyperglycemia, of *Apc^Min/+^* mice (**Fig. 6E-F** and **S9A-B**). We also observed improvements in the loss of gastrocnemius (Gas) and tibialis anterior (TA) muscle and inguinal (iWAT) and epididymal (e-WAT) white adipose tissues, as well as decreased myofiber cross-sectional area and adipocyte size, of *Apc^Min/+^* mice (**Fig. 6G-H** and **S10A**). We used another GCGR inhibitor, LGD-6972 (LGD), to avoid off-target effects and consistently observed the improvement of tumor-induced wasting in *Apc^Min/+^* mice without affecting tumor growth (**Fig. S9C-H** and **S10B**).

We also studied mice bearing Lewis Lung Carcinoma (LLC) tumors, an established lung-cancer-cachexia mouse model. Similar to *Apc^Min/+^* mice, we observed hyperglucagonemia, enlarged α-cell mass, weight decline, and loss of muscle and fat tissues in *C57BL/6* mice bearing LLC tumors at 21 days after LLC injection (**Fig. 6I-M, S9I-K** and **S10C**). Injection of LLC cells in *Gcg^-/-^* mice, as compared to *C57BL/6* mice, did not significantly affect tumor growth but, strikingly, alleviated host wasting including weight decline, hyperglycemia, and loss of fat and muscle (**Fig. 6J-M, S9I-K** and **S10C**). In addition, we IP injected GRA into LLC-tumor bearing mice to blunt glucagon response for 7 days and observed no effects on tumor growth but significant improvements in weight decline, hyperglycemia, loss of muscle and fat, and hepatic glycogen depletion (**Fig. 6J-M, S9I-K** and **S10C**). Taken together, our results demonstrate that malignant tumors cause organ wasting via elevation of glucagon production.

### PDGFR/VEGFR blockade alleviates cholinergic-α-cell contacts, glucagon production and wasting in tumor-bearing mice

Similar to Akh secretion modulated by cholinergic neurons, mammalian glucagon release is controlled by acetylcholine probably derived from parasympathetic nerves^56^. Published single-cell RNA-seq data reveal that α-cells express acetylcholine receptors as well as VEGFRs and PDGFRs (VEGFR/PDGFR), the homolog of *Drosophila* Pvr ^57,58^ (**Fig. S10E**). Because mouse *Apc^Min/+^* colon, LLC lung, and human pancreatic tumors were found to produce large amounts of VEGF/PDGF ^59–66^, we wonder whether tumors promote neural-associated glucagon secretion via VEGFR/PDGFR signaling in a manner similar to fly Pvr signaling. To address this hypothesis, we treated cultured glucagon-producing αTC1 cells with synthetic PDGF-BB to activate PDGFR/VEGFR signaling and observed an increase in expression of multiple *Mmp* genes and a decrease in *Timp* genes (**Fig. S10F**). In line with this, we next examined islet morphologies in *Apc^Min/+^* mice and found less ECM contents in α-cells that were indicated by versican staining (**Fig. 7A**). Meanwhile, we strikingly observed increased overlapping between α-cells and intrapancreatic nerves (PGP9.5^+^), especially nerves (AchT^+^), in *Apc^Min/+^* mice, indicating cholinergic innervation of α-cells (**Fig. 7B** and **S10G**). The cholinergic innervation was further associated with the regions of α-cell expansion (**Fig. 7B**).

**Figure 7.**
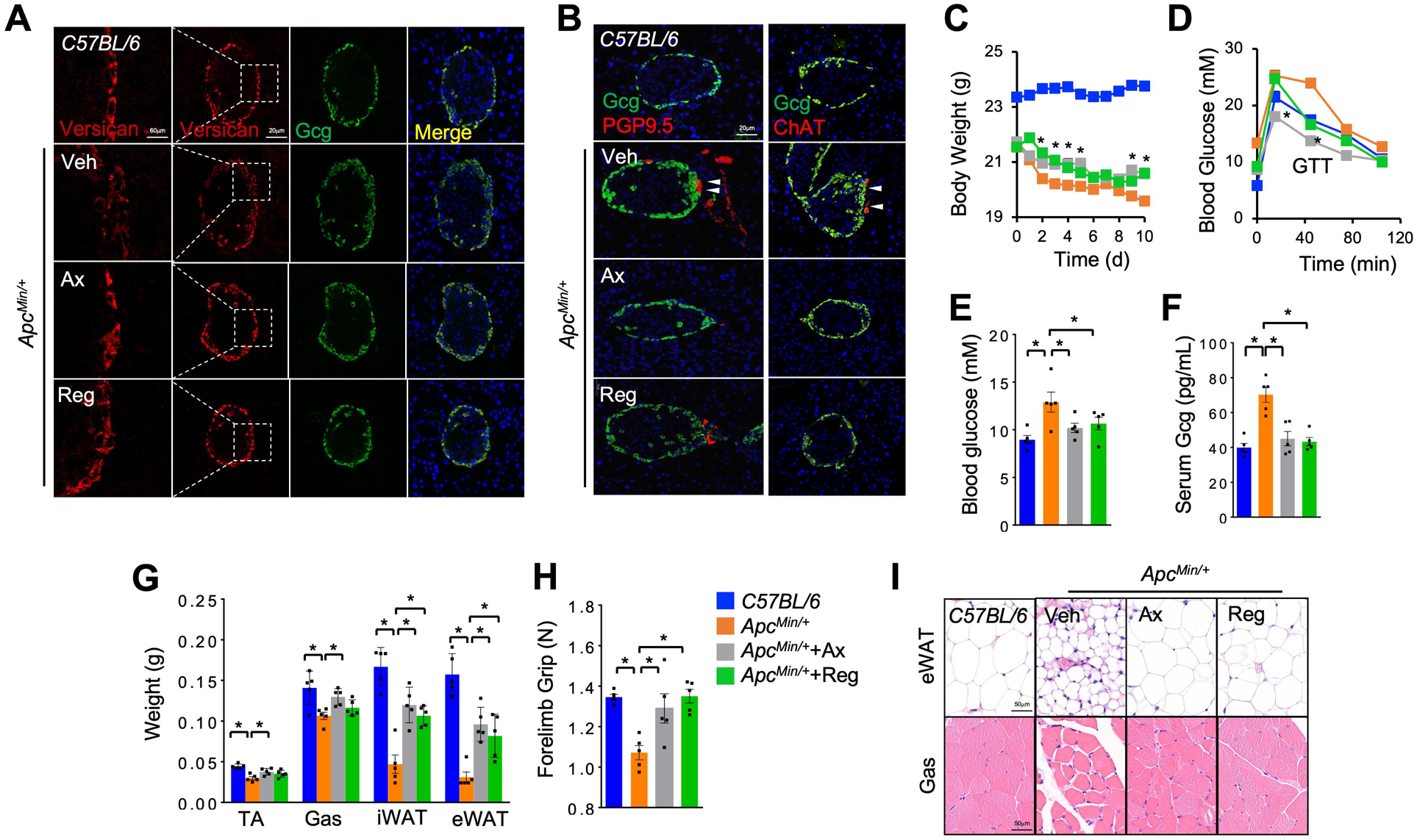
PDGFR/VEGFR blockade alleviates hyperglucagonemia and cholinergic-α-cell contacts in tumor-bearing mice. (**A-I**) ECM levels of α- cells (**A**, green, anti-Gcg; red, anti-Versican) and islet morphologies (**B**, green, Gcg; left, red, anti-PGP9.5; right, red, anti-ChAT) indicated by confocal images, serum glucagon levels (**C**), body weights (**D**), GTT (**E**), fed blood glucose (**F**), tissue weights (**G**), forelimb grip strength (**H**), and tissue morphologies (**I**, Gas, myotube diameters; eWAT, adipocyte sizes) of indicated mice that were performed daily IP injection of PDGFR/VEGFR inhibitors from week 16 for 2 weeks. *C57BL/6* mice (control), *Apc^Min^*+veh, *Apc^Min^*+Ax (30 mg/Kg/day), and *Apc^Min^*+Reg (20 mg/Kg/day) (n=5). Data are presented as mean ± SEM. Each dot represents one biological replicate. Statistical analysis was conducted by one-way ANOVA with Bonferroni’s multiple-comparisons test. \**p* < 0.05.

To investigate the impacts of PDGFR/VEGFR signaling on cholinergic innervations and glucagon secretion, we IP injected VEGFR/PDGFR inhibitors, axitinib (Ax) or regorafenib (Reg), into *Apc^Min/+^* mice to block VEGFR/PDGFR signaling at week 18. We found that short-term administration of either inhibitor robustly restored ECM contents of α-cells and diminished cholinergic innervation and α-cell expansion to alleviate hyperglucagonemia (**Fig. 7A-B**). In support of glucagon catabolic effects, we subsequently found that either Ax or Reg administration hardly affected tumor growth but significantly improved the weight loss, glucose intolerance, loss of fat and muscle tissues, and forelimb weakness in *Apc^Min/+^* mice (**Fig. 7C-I, S10D** and **S10H**). Collectively, these results demonstrate that PDGFR/VEGFR blockade decreases cholinergic innervation on α-cells, glucagon release, and host wasting in *Apc^Min/+^* mice.

## Discussion

Tumor-induced host organ wasting is a general phenomenon in both vertebrates and invertebrates. We have previously revealed that yki^3SA^-gut tumors cause host wasting partially through secretion of ImpL2 and Upd3 to impair local metabolism of muscle or fat tissue using *Drosophila* as a conserved cancer model ^9,10^. In this study, we further uncovered that yki^3SA^ tumors extensively result in systemic energy wasting via secretion of another ligand Pvf1 to hijack neuronal-associated release of Akh, a critical metabolic hormone from host. The molecular mechanisms include that Pvf1-Pvr axis triggers ECM remodeling of APCs and enhances their innervation of excitatory cholinergic neurons (**Fig. 5I**). We also identified the similar neural regulation of glucagon production regarding organ wasting in tumor-bearing mice.

It is interesting to find that Akh deficiency robustly improved muscle dysfunction and ovary atrophy, in addition to established lipid loss and hyperglycemia, in tumor-bearing flies, even though *AkhR* is not detectable in either muscle or ovary cells (FlyCellAtlas ^35^). We speculate an indirect regulation of muscle and ovary homeostasis by Akh. Note that, AkhR removal in both fat body and neurons partially alleviated host wasting. It could be possible that AkhR signaling in the fat body causes systemic amino acid consumption ^25^, leading to and subsequent ovary and muscle degeneration. AkhR^+^ neurons might promote muscle or ovary wasting through directly or indirectly projections in response to excessive Akh as well. However, Akh gain-of-function in control non-tumor flies failed to affect organ wasting, such as muscle dysfunction and ovary atrophy, suggesting that Akh might collaborate with other tumor-associated factors like inflammatory responses to regulate organ wasting in *yki3SA* tumor-bearing flies.

We have previously characterized the catabolic roles of tumor-derived Pvf1 and speculated that fat body and muscle as the major responding tissues based on Pvr gain-of-function in non-tumor flies ^11^. However, the real functional target organ(s) of Pvf1 validated by Pvr loss-of-function in the yki^3SA^-tumor-bearing flies are still unclear. In this study, we demonstrated APCs, but not adipose or muscle, as the predominant tissues that respond to tumor-derived Pvf1 to cause wasting. This is because Pvr inactivation in either muscle or fat body in flies bearing Pvf1-inducing tumors rarely shows wasting improvement. The discrepancies of Pvr effects between APCs and fat body/muscle would be caused by differential *in vivo* Pvf1 delivery, abundance of Pvr expression, and/or intratissue signaling regulation in yki^3SA^-tumor-bearing flies. In line with our speculations, recent evidence showed that Pvf1-Pvr signaling in Malpighian tubules contributes to yki^3SA^-tumor-induced organ wasting. It would be interesting to investigate the impacts of Pvr signaling in additional tissues like oenocytes ^67^ and neurons of yki^3SA^-tumor-bearing flies using *LexA/GAL4* binary expression system in future studies.

Pvf1-Pvr activation in APCs increases both mRNA level and release of Akh. We investigated Pvr downstream regulators and found that *ERK* knockdown restores both of them, while *Mmp2* knockdown in APCs only alleviates Akh release without affecting *Akh* transcription, in the context of Pvf1-Pvr activation. These results suggest that Pvf1-Pvr axis in APCs promotes Akh release via not only ERK-associated Akh synthesis, but also ERK-Mmp2-induced Akh secretion. Consistent with this notion, our results further revealed that tumor-derived Pvf1 activates Pvr in APCs to increase Mmp2-dependent ECM degradation and APC innervation to upstream neurons, promoting Akh release.

Previous studies have reported that upstream inhibitory neurons project to APCs to suppress Akh release in response to nutrient availability ^30–32^, however, the existence of upstream excitatory neurons of APCs to promote Akh release in *Drosophila* was largely unknown. By screening multiple neurotransmitters in this study, we identified cholinergic neurons as the major excitatory neurons innervating APCs. Overexpression of TrpA1 and Shi^TS^ to activate cholinergic neurons and impair the projection to APCs, respectively, further confirmed their pivotal roles in promoting Akh release and enhancing systemic energy wasting in the context of yki^3SA^ tumors. Because multiple acetylcholine receptors were found to be expressed in APCs, cholinergic neurons most likely release acetylcholine to directly impact on APC function. In addition, as compared to cholinergic neurons, we also observed relatively weaker contacts between APCs and neurons labelled by *TRH-*, *Gad1-*, and *Tdc2-GAL4*. A recent study also indicated that *Tdc2-GAL4-*labelled octopaminergic neurons project on APCs ^68^. The potential of contacts between APCs and these neurons might impact Akh release as well in normal or yki^3SA^-tumor-bearing flies. On the other hand, the single-nucleus RNA-seq that indicates moderate *Pvr* expression in, at least, cholinergic and serotonergic neurons (FlyCellAtlas.org ^35^) raises the possibility that Pvf1 might also modulate the activity of these upstream innervating neurons to coordinate Akh production.

One of our important findings here includes that glucagon is essential for muscle and fat wasting in tumor-bearing mice. The plausible mechanisms include at least glucagon-associated hepatic amino acid catabolism remotely causes systemic amino acid loss and organ wasting ^54,69^. Taken together previous studies that demonstrate glucagon’s roles in systemic wasting regulation in type 2 diabetic and sleep-deprived mice ^29,54^, we propose that glucagon also functions as a general cachectic hormone in chronic conditions, beyond its established role in short-term hyperglycemic regulation. While anorexia resulted from malignant cancer invariably increases glucagon release that potentially contribute to energy wasting ^70^, elucidating the pathogenic mechanisms that stimulate glucagon production by critical tumor-associated cachectic factors remains a crucial research endeavor.

*Ex vivo* studies have revealed that multiple mammalian neurotransmitters including acetylcholine control glucagon secretion ^56,71^. However, there are very limited *in vivo* evidence. Thus, it is striking to observe cholinergic-α-cell contact in the context of malignant tumors. We further revealed the VEGFR/PDGFR signaling promotes cholinergic-α-cell interaction through ECM remodeling of α-cells in tumor-bearing *Apc^Min/+^* mice, leading to excessive glucagon release and host wasting. Even though we observed PDGF-BB-induced MMPs expression in cultured α-TC1 cells, incorporating both α-cells and cholinergic neurons into organoids would help provide a more definitive confirmation of neural-associated glucagon release in α-cells in the future. Note that, beside glucagon secretion, α-cell proliferation is also correlated to cholinergic contact with unknown mechanisms. Given large amounts of VEGF/PDGFs produced in both rodent and human malignant tumors ^59–62^, we therefore conclude a novel mechanism of tumor-host interaction whereby tumors remotely enhance cholinergic-α-cell innervation via VEGFR/PDGFR signaling to promote glucagon release and systemic energy loss.

## Author Contributions

G.D., Y.L. and C.C. designed and performed experiments, including metabolic assays, qPCR, genetic manipulation, and neuronal analysis. G.D., Y.L., C.C. and K.T. performed mouse work. Y.D., H.P., Z.W., P.D., E. R., and X.W. helped perform mouse work and fly genetics. Y.Y., Z. Y. and W.S. discussed results. C.C. and W.S. wrote the manuscript.

## Acknowledgements

We thank the DRSC/TRiP at Harvard Medical School, Bloomington Drosophila Stock Center, NIG in Japan, and Vienna Drosophila Resource Center for providing fly stocks; Sheng Li (South China Normal University) for *Vkg-GFP* and *UAS-Timp* flies and integrin βPS antibodies; Ronald Kühnlein (Max-Planck Institute) for *Akh^SAP^ and Akh^A^*flies; Norbert Perrimon (Harvard Medical School) for *Akh-GAL4* flies; Wei Zhang (Tsinghua University) for *Cha-GAL4;* Yufeng Pan (Southeast University, China) for *Th-GAL4, Tdc2-GAL4, Trh-GAL4* and *Oct-TyrR-GAL4* lines; Yong Liu and Baoliang Song (Wuhan University, China) and Liangyou Rui (Michigan University) for insightful comments. Work in the Song lab was supported by the Chinese National Natural Science Foundation (92357303, 91957118, 32350013, 32425029), Chinese Ministry of Science and Technology (National Key R&D Program, 2021YFC2700700) and the Fundamental Research Funds for the Central Universities (2042022dx0003).

## Supplemental Figure Legends

**Supplemental Figure 1. Akh is essential for tumor-induced host wasting in *Drosophila*.** (**A-B**) Representative images of muscle degeneration (**A,** up) indicated by swollen mitochondria (M) and gaps (G) between mitochondria and myofibril (F), ovary atrophy (**A,** down), and climbing rates (**B,** n=17) of adult yki^3SA^-tumor-bearing flies with or without *Akh^Δ^* mutation (*Akh^SAP^/Akh^A^*) at day 8. (**C-D**) Metabolic changes including TAG and trehalose (TRE) storages in 8-day old indicated control adult flies (**C,** n=5; **D,** n=6). (**E**) PCA analysis of whole-body gene expression of yki^3SA^-tumor-bearing flies with or without *Akh^Δ^* mutation (*Akh^SAP^/Akh^A^*) at day 8. (**F**) Gene Ontology Enrichment analysis of 395 differentially expressed genes that are both yki^3SA^-tumor- and Akh-dependent indicating that the following terms of biological process, cellular compartment, and KEGG pathways are significantly enriched. Data are presented as mean ± SEM. Each dot represents one biological replicate. Statistical analysis was conducted by two-tailed unpaired t-test (**C**) or one-way ANOVA with Bonferroni’s multiple-comparisons test (**B, D**). \**p* < 0.05.

**Supplemental Figure 2. AkhR in either fat body or brain contributes to tumor-induced host wasting.** Representative images of abdomen bloating (**A, F,** up), gut tumors and mass quantification (**A, F,** middle, GFP; **B, G,** n=3), ovary atrophy (**A, F,** bottom), bloating rates (**C, H,** n=3), climbing rates (**D, I,** n=20), and global storages of TAG and trehalose (TRE) (**E, J,** n=6) of adult yki^3SA^ tumor-bearing flies (LexA+*GAL4*) with *AkhR* RNAi in either fat body (*R4-GAL4>*) (**A-E**) or pan-neuron (*elav-GAL4>*) (**F-J**) at day 4. Data are presented as mean ± SEM. Each dot represents one biological replicate. Statistical analysis was conducted by one-way ANOVA with Bonferroni’s multiple-comparisons test. \**p* < 0.05.

**Supplemental Figure 3. Excessive Akh release causes lipid and carbohydrate mobilization but not organ wasting.** Representative images of abdomen (**A, F, K, N**), whole-body gene expression (**B, G,** n=4, 5 flies/replicate), whole-body TAG and TRE levels (**C, H,** n=4, 5 flies/replicate), climbing rates (**D, I, L, O,** n=20), as well as ovary images and size quantification (**E, J, M, P,** n>10), of adult flies with TrpA1 overexpression in the APCs (**A-E**), Akh overexpression in the APCs (*Akh-GAL4>,* **F-J**), fat body (*R4-GAL4,* **K-M**), or pan-neurons (*elav-GAL4,* **N-P**) at 29 degree at day 8 (**A-J**) or 25 degree at day 4 (**K-P**). Data are presented as mean ± SEM. Each dot represents one biological replicate. Statistical analysis was conducted by two-tailed unpaired t-test. \**p* < 0.05.

**Supplemental Figure 4. Pvr autonomously regulates Akh production.** (**A**) RNAi against indicated genes that affect larval glycemic levels by >10% in APCs. (**B**) Immunostaining indicating Pvr expression in the larval APCs (green, Pvr>GFP; red, anti-Akh). (**C**) Glycemic (circulating trehalose) levels of indicated flies (left, larvae at day 5, n=8, 10 flies/replicate; right, adult flies at day 4, n=3, 30 flies/replicate). (**D**) Representative images of CalexA.GFP indicating intracellular Ca^2+^ flux in adult flies with Pvr^AC^ overexpression in the APCs (left) and GFP quantification (right, n=6). (**E-H**) Circulating Akh levels (**E,** left, dot-blot; right, quantification, n=3), systemic mRNA levels of *Akh* and *tobi* (**F**, n=3, 5 flies/replicates), TAG and trehalose (TRE) storages (**G**, n=4, 5 flies/replicates), and survival under starvation (**H,** n=4, 20 flies/replicate) of indicated flies at day 4. (**I-L**) Circulating Akh levels (**I,** dot-blot), Akh production in APCs (**J**, left) and quantification of APC masses and intracellular Akh amounts (**J,** right, n=3), TAG and trehalose (TRE) (**K**, n=4, 5 flies/replicates), and survival under starvation (**L,** n=4, 20 flies/replicate) of indicated flies with Pvr manipulation only in adult APCs using *tub-Gal80^TS^*at day 5 after transgene induction. Data are presented as mean ± SEM. Each dot represents one biological replicate. Statistical analysis was conducted by two-tailed unpaired t-test (**D-G, J-K**) or log-rank test (**H, L**). \**p* < 0.05.

**Supplemental Figure 5. Pvr blockade in APCs alleviates tumor-induced wasting.** Representative images of abdomen bloating (**C,** up), gut tumors and mass quantification (**C,** middle, GFP; **D,** n=3), ovary atrophy (**A,** up; **C,** middle) and muscle degeneration (**A, C,** bottom) indicated by swollen mitochondria (M) and gaps (G) between mitochondria and myofibril (F), climbing rates (**B,** n=17; **F,** n=17), bloating rates (**E,** n=4), and metabolic changes including TAG and trehalose (TRE) storages (**G,** n=4) of yki^3SA^-tumor-bearing flies (*LexA+GAL4*) with APC Pvr inactivation (Pvr^DN^, **A-B**) or Pvr RNAi (**C-G**) at day 6. Data are presented as mean ± SEM. Each dot represents one biological replicate. Statistical analysis was conducted by one-way ANOVA with Bonferroni’s multiple-comparisons test. \**p* < 0.05.

**Supplemental Figure 6. Pvr blockade in muscle or fat body hardly alleviates tumor-induced wasting.** Representative images of abdomen bloating (**A, F,** up), gut tumors and mass quantification (**A, F,** middle, GFP; **B, G**, n=3), muscle degeneration (**A, F,** bottom) indicated by swollen mitochondria (M) and gaps (G) between mitochondria and myofibril (F), bloating rates (**C,** n=4; **H,** n=3), climbing rates (**D,** n=17; **I,** n=20), and metabolic changes including TAG and trehalose (TRE) storages (**E,** n=4; **J,** n=6) of yki^3SA^-tumor-bearing flies (*LexA+GAL4*) with Pvr inactivation (Pvr^DN^) in the fat body (*R4>,* **A-E**) or muscle *(Mhc>,* **F-J**) at day 6. Data are presented as mean ± SEM. Each dot represents one biological replicate. Statistical analysis was conducted by one-way ANOVA with Bonferroni’s multiple-comparisons test. \**p* < 0.05.

**Supplemental Figure 7. Pvr regulation of Akh production.** Survival under starvation (**A, E, H, L,** n=4, 20 flies/replicate), whole body *Akh* or *tobi* expression (**B, D, J,** n=3 or 5, 5 flies/replicate), metabolic changes such as TAG or TRE storages (**C, F, I, K,** n=3 or 5, 5 flies/replicate) and circulating Akh levels in the hemolymph (**G**) of adult flies with indicated genotypes at day 4 (**A-I**) or 8 (**J-L**). Data are presented as mean ± SEM. Each dot represents one biological replicate. Statistical analysis was conducted by two-tailed unpaired t-test (**B-D, F, I, J,** left, **K,** left), one-way ANOVA with Bonferroni’s multiple-comparisons test (**J,** righ**, K,** right), or log-rank test (**A, E, H, L**). \**p* < 0.05.

**Supplemental Figure 8. ECM homeostasis and neural innervation of APCs.** (**A-B**) Representative images of ECM homeostasis indicated by Integrin βPS (**A, B,** left, red, anti-integrin) around the somas of APCs (**A**, left, green, GFP; **B,** left, green, anti-Akh) and quantification of extracellular integrin amounts (**A, B,** n=5-9) of adult flies with Pvr^AC^ overexpression in APCs at day 4 or flies bearing yki^3SA^-tumor plus *Pvf1 R*NAi at day 7. (**C**) Representative images of boutons that are indicated by Syt-GFP (green) driven by indicated GAL4 lines in the somas of adult APCs (red, Akh) at day 4. (**D**) Published snRNA-seq data indicating that multiple neurotransmitter receptors are expressed in adult CCs or APCs. Data are presented as mean ± SEM. Each dot represents one biological replicate. Statistical analysis was conducted by two-tailed unpaired t-test (**A**) or one-way ANOVA with Bonferroni’s multiple-comparisons test (**B**). \**p* < 0.05.

**Supplemental Figure 9. Blockade of glucagon response alleviates wasting in tumor-bearing mice.** (**A-B**) Hepatic gene expression (**A**) and glucose tolerance test (GTT) (**B**) of *Apc^Min/+^* mice from week 18 with or without daily IP injection of 10 mg/Kg/day GRA Ex-25 (GRA) for two weeks (n=10). (**C-H**) Gut tumors (**C**), body weights (**D**), fed blood glucose (**E**), tissue weights (**F**), forelimb grip strengths (**G**), and tissue morphologies (**H**, Gas, myotube diameters; eWAT, adipocyte sizes) of *Apc^Min/+^* mice from week 18 with or without IP injection of 6 mg/Kg/day LGD-6972 (LGD) for 10 days (two injections in every three days) for two weeks (n=5). (**I-K**) Fed blood glucose levels (**I**), tissue weights (**J**), and liver glycogen contents (**K**, PAS staining) of indicated LLC-tumor-bearing mice (*C57BL/6*, n=5; *C57BL/6*+LLC, n=6; *Gcg^-/-^ +*LLC, n=10) or LLC-tumor-bearing mice with daily IP injection of GRA from day 14 for 7 days (PBS, n=6; LLC, n=7; LLC+GRA, n=8, 10 mg/Kg/day). Data are presented as mean ± SEM. Each dot represents one biological replicate. Statistical analysis was conducted by two-tailed unpaired t-test (**A-B**) or one-way ANOVA with Bonferroni’s multiple-comparisons test (**D-G, I-J**). \**p* < 0.05.

**Supplemental Figure 10. PDGFR/VEGFR express in α-cells.** (**A-D**) Quantification of gastrocnemius myofiber cross-sectional area, epididymal white adipocyte size, as well as hepatic glycogen content (PAS intensity) in the indicated mice (n=5). (**E**) Gene expression in both mouse α- and β-cells as indicated by RPKM in a published dataset (GSE54973). Genes with RPKM > 1 are considered as functional expressed. (**F**) Gene expression in α- TC1 cells that were treated with 10 ng/mL PDGF-BB for 3 hours (n=3). (**G**) Surface rendering of confocal images merging 8 sections of neve-α-cell contact the islet from the top side (green, Gcg; red, PGP9.5). (**H**) Tumor morphologies indicated by Methylene-blue staining in the colon (arrows indicate tumors) of that were performed daily IP injection of PDGFR/VEGFR inhibitors from week 16 for 2 weeks. *C57BL/6* mice (control), *Apc^Min^*+Veh, *Apc^Min^*+Ax (30 mg/Kg/day), and *Apc^Min^*+Reg (20 mg/Kg/day) (n=5). Data are presented as mean ± SEM. Each dot represents one biological replicate. Statistical analysis was conducted by two-tailed unpaired t-test (**F**) or one-way ANOVA with Bonferroni’s multiple-comparisons test (**A-D**). \**p* < 0.05.

## Supplemental Table Legends

**Supplemental Table 1. The FPKM values of gene expression in the whole body of yki^3SA^-tumor-bearing flies with or without Akh mutation.**

**Supplemental Table 2. The glycemic changes of larvae with APCs bearing different RNAi against transmembrane proteins.**

**Supplemental Table 3. Clinical characteristics of patients with pancreatic cancer and benign diseases.**

## Experimental Methods

### Fly strains

Files were raised on fly food (5 g agar, 25 g dry yeast, 75 g corn flour, 90 g sucrose, 1.5 g Methylparaben, 4 mL propionic acid per liter) in incubator 12 h light /12 h dark cycle at 25°C.

Transgenic and mutant flies were used as below:

| Genotypes | Source | Identifier |
| --- | --- | --- |
| *esg-GAL4, tub-GAL80<sup>TS</sup>, UAS-GFP* | <sup>1</sup> | N/A |
| *R4-Gal4* | <sup>1</sup> | N/A |
| *Mhc-Gal4* | <sup>1</sup> | N/A |
| *elav-Gal4* | <sup>2</sup> | N/A |
| *w<sup>1118</sup>* | <sup>1</sup> | N/A |
| *UAS-w-RNAi* | <sup>1</sup> | N/A |
| *UAS-Raf<sup>F179</sup>* | <sup>1</sup> | N/A |
| *Vkg-GFP* | A gift from Dr. Sheng Li (South China Normal University) <sup>3</sup> | N/A |
| *Akh<sup>SAP</sup>* | A gift from Dr. Ronald Kühnlein (Max-Planck Institute) <sup>4</sup> | N/A |
| *Akh<sup>A</sup>* | A gift from Dr. Ronald Kühnlein (Max-Planck Institute) <sup>4</sup> | N/A |
| *UAS-Timp* | Bloomington Stock Center | BDSC_58708 |
| *Pvr-Gal4* | Bloomington Stock Center | BDSC_76657 |
| *Cha-GAL4* | Bloomington Stock Center | BDSC_6798 |
| *UAS-shi<sup>TS</sup>* | Bloomington Stock Center | BDSC_44222 |
| *Trh-GAL4* | Bloomington Stock Center | BDSC_38388 |
| *Vglut-GAL4* | Bloomington Stock Center | BDSC_60312 |
| *Gad1-GAL4* | Bloomington Stock Center | BDSC_51630 |
| *Th-GAL4* | Bloomington Stock Center | BDSC_8848 |
| *Tdc2-GAL4* | Bloomington Stock Center | BDSC_9313 |
| *Akh-GAL4* | Bloomington Stock Center | BDSC_25684 |
| *UAS-Denmark* | Bloomington Stock Center | BDSC_33062 |
| *UAS-Akh* | Bloomington Stock Center | BDSC_27343 |
| *UAS-TrpA1* | Bloomington Stock Center | BDSC_26263 |
| *UAS-Syt-GFP* | Bloomington Stock Center | BDSC_6926 |
| *UAS-Pvr-RNAi* | Bloomington Stock Center | BDSC_37520 |
| *UAS-Pvr<sup>AC</sup>* | Bloomington Stock Center | BDSC_58428 |
| *UAS-Pvr<sup>DN</sup>* | Bloomington Stock Center | BDSC_58431 |
| *UAS-Mmp1-RNAi1* | Vienna Drosophila Resource Center | v101505 |
| *UAS-Mmp1-RNAi2* | Bloomington Stock Center | BDSC_31489 |
| *UAS-Mmp2-RNAi1* | National Institute of Genetics | NIG_1794-1R-1 |
| *UAS-Mmp2-RNAi2* | Bloomington Stock Center | BDSC_61309 |
| *UAS-ERK-RNAi* | Vienna Drosophila Resource Center | v109108 |
| *UAS-Pvf1-RNAi* | Vienna Drosophila Resource Center | v102699 |
| *UAS-Akhr-RNAi* | Bloomington Stock Center | BDSC_29577 |
| *esg-LexA.RJ-4* | Bloomington Stock Center | BDSC_66630 |
| *LexAop-GFP* | Bloomington Stock Center | BDSC_32205 |
| *LexAop-cdTom* | Bloomington Stock Center | BDSC_77139 |
| *UAS-CaLexA.GFP* | Bloomington Stock Center | BDSC_66542 |
| *Trans-tango* | Bloomington Stock Center | BDSC_77124 |
| *UAS-yki<sup>3SA</sup>* | Bloomington Stock Center | BDSC_28817 |
| *LexAop-yki<sup>3SA</sup> (attP40)* | This study | N/A |
| *LexAop-yki<sup>3SA</sup> (attP2)* | This study | N/A |

To induce gut tumors, we followed the experimental procedures described previously ^1^. Briefly, different *UAS* and *LexAop* insertions were crossed to *esg-GAL4, tub-GAL80^TS^, UAS-GFP* and *esg-LexA, tub-GAL80^TS^, LexAop-GFP* at 18°C, respectively, to inactivate GAL4/LexA. 4-day-old virgin adult progenies were placed at 29°C to induce the transgenes (day 0 for tumor induction). Flies were transferred onto fresh food every 2 days.

For the *in vivo* RNAi screening against transmembrane proteins in larval APCs, different RNAi lines from DRSC/TRiP center of Harvard Medical School were crossed to *Akh-GAL4* at 25 °C. The off-spring larvae were allowed to grow at 25 °C for 5 days to reach late 3^rd^-instar for glycemic measurements as previously reported ^5^. To access the regulation of Akh release in adult flies, different UAS lines were crossed to *Akh-GAL4* at 25 °C. The virgin progenies were collected and maintained at 25 °C for 4 days for metabolic and biochemical measurements. Negative controls, *w^1118^* and *UAS-w-RNAi,* exhibited similar phenotypes and only *w^1118^* is shown in the figures.

The 3^rd^ instar larvae and female adult flies were used for metabolic and wasting examination in this study.

### Lipid and carbohydrate measurements in flies

We measured fly TAG and carbohydrates as described previously ^1^. Briefly, 5 female flies from each group were homogenized with 0.5 mL PBS containing 0.2% Triton X-100 using Multi-sample tissuelyser-24 (Shanghai Jingxin Technology) and heated at 70°C for 5 min. The supernatant was collected after centrifugation at 12,000 X g for 10 min at 4°C. 10 μL of supernatant was used for protein quantification using Bradford Reagent (Sigma, B6916- 500ML). Whole body trehalose levels were measured from 10 μL of supernatant treated with 0.2 μL trehalase (Megazyme, E-TREH) at 37°C for 30 min using glucose assay reagent (Megazyme, K-GLUC) following the manufacturer’s protocol. We subtracted the amount of free glucose from the measurement and then normalized the subtracted values to protein levels in the supernatant. To measure whole body triglyceride levels, we processed 10 μL of supernatant using a Serum Triglyceride Determination kit (Sigma, TR0100), subtracted the amount of free glycerol in the supernatant from the measurement, and then normalized to protein levels in the supernatant.

### Dot-blot analysis of hemolymph Akh

Hemolymph of 60–80 female adult flies was collected and 1:100 diluted with PBS. 10 μL of diluted hemolymph was dropped on nitrocellulose membrane (GE Healthcare) and air dried at room temperature for 5 min. The membrane was then boiled in PBS for 3 min and subsequently fixed with 4% PFA in PBS for 20 min. The membrane was blocked with 3% BSA in PBS for 30 min at room temperature and then incubated with rabbit anti-Akh antibody (1:1000, Abclonal, A22867) in 3% BSA in PBS at 4 °C for overnight followed by incubation with HRP-conjugated secondary antibodies in 3% BSA in PBS for 1 h at room temperature. The intensities of the black dots were considered as the amounts of Akh in the hemolymph. Ponceau Red staining prior to blocking was used as the loading control.

### Climbing, food intake and starvation assays

Female adult flies, which were placed in an empty vial and tapped down to the bottom for climbing, were allowed to climb for 2 s. Flies were filmed to quantify climbing height and speed using ImageJ. A minimum of 15 flies and 3 independent trials were performed for each condition. 5 female adult flies, which were cultured on normal food containing 1% (w/v) bright-blue dye for 24h, were homogenized in 1 mL PBS and centrifuged at 12,000 X g for 10 min at 4°C to remove the debris. Protein concentration in the 10 μL lysate was measured using Bradford Reagent (Sigma, B6916-500ML). The food intake was measured by quantifying concentration of bright-blue dye in 200 μL lysate at 595 nm with normalization to protein concentration. 80 female adult flies of genotype were cultured on starvation food (1% agar in H_2_O). Dead flies were counted every 12 h.

### Mice strains and cell lines

All mouse work was approved by the Animal Care and Ethical Committee at Wuhan University. *C57BL/6, Apc^Min/+^* (T001457), and *Gcg^-/-^* (T014382) mice were obtained from Gempharmatech, China and housed individually at 22-24°C with a 12 h light/dark cycle with water and food ad libitum. Sample size, determined empirically via performing preliminary experiments, was chosen to be at least five to ensure that adequate statistical power was achieved. Male mice were grouped with similar average body weight one week prior for inhibitor administration. Tumor-bearing mice received daily intraperitoneal injections of Axitinib (T1452, Targetmol) (30 mg/Kg/day), Regorafenib (T1792, Targetmol) (20 mg/Kg/day), GRA-Ex-25 (T3422, Targetmol) (10 mg/Kg/day) or vehicle from 16-18 weeks. LGD-6972 (T25711, Targetmol) (6 mg/Kg/day, two injections every three days) or vehicle was intraperitoneally injected into mice from 18-20 weeks. As for LLC-tumor-bearing mice, PBS or 5 million LLC cells per mouse were injected subcutaneously over the flank (Day 0). After LLC- cell progression for 14 days (Day 14), mice received daily intraperitoneal injections of vehicle or 10 mg/kg GRA-Ex-25 (T3422, Targetmol) for 7 days (Day 21).

All mice were sacrificed in the late light cycle (3 p.m.–6 p.m.) and tissues were weighed and frozen immediately in liquid nitrogen for future tests.

αTC1 cells were cultured in DMEM (PM150211, ProCell, China) supplemented with 1g/L glucose, 10% FBS and antibiotics. After incubation with 10 ng/mL PDGF-BB (AP002631HU, Cusabio, China) for 3 hours, αTC1 cells were washed and lysed using Trizol (15596018, Thermo Fisher) for RNA extraction.

### Data of patients with pancreatic cancer

The study was performed in accordance with the Helsinki Declaration and approved by the Medical Ethics Committee of Zhongnan Hospital of Wuhan University (KELUN2021042). All participants gave their written informed consent. Patient-, surgery- and oncology-related data were obtained from medical records. Weight loss > 1.5% per month within three months before surgery was considered as weight decline. Peripheral blood was drawn from patients with pancreatic cancer and benign diseases who were diagnosed by pathological examination at Department of Hepatobiliary and Pancreatic Surgery of Zhongnan Hospital of Wuhan University. The plasma was separated and stored at −80 °C until analysis.

### Immunostaining and electronic microscopy

Adult midguts, larval and adult brains were dissected in PBS and fixed for 15 min in PBS containing 4% paraformaldehyde. After fixation, the samples were washed with PBST (0.2% Triton-X100 in PBS) and blocked with 1% BSA in PBST. After incubation with primary antibodies anti-Akh (1:10000, this study) or integrin βPS (1:100, DSHB, CF.6G11) overnight at 4°C, the tissues were washed and then incubated with Alexa fluorescence secondary antibody (1:1000, A32742, Thermo Fisher) and DAPI (1:1000, D1306, ThermoFisher) for 1 h at room temperature then washed. Images of fly appearances were performed on a Nikon SMZ18 or Nikon Eclipse Ts2 and confocal images were obtained using a Zeiss LSM880. For quantification of Akh staining and CaLexA-GFP intensities, stacks were Z-projected and the signals of and Akh or CaLexA-GFP in the whole CC were quantified using Integrated Density in ImageJ. The background was subtracted to give the net signal. APC masses were quantified using ImageJ to analyze the APC areas.

Thoraces from adult flies were fixed in 0.1 M sodium cacodylate buffer (pH 7.4) containing 2.5% glutaraldehyde, 2% paraformaldehyde overnight and were performed EM analysis following standard protocols by Servicebio at Wuhan, China.

Mouse and human pancreatic samples were fixed in PBS containing 4% paraformaldehyde and cyrosections were performed following standard protocols by Servicebio at Wuhan, China. Primary antibodies including anti-glucagon (1:500, Servicebio, GB12097), anti-PGP9.5 (1:200, Servicebio, GB11159-1), anti-Versican (6 mg/mL, Sigma, AB1033), biotinylated HABP (1:500, Sigma, 385911); Alexa fluorescence secondary antibodies (1:300, Servicebio, GB21303) and (1:400, Servicebio, GB25301) together with DAPI (1:1000, Servicebio, G1012-100ML) and HRP-labeled Streptavidin (1:2000, Beyotime, A0303) were used. Confocal images were obtained using a Zeiss LSM880. Confocal images of 8 sections of a pancreas/islet were merged to generate the 3D-view image.

Adipose, liver, and gut tissues were fixed in PBS containing 4% paraformaldehyde and were performed H&E, PAS, or Methylene-blue staining following standard protocols by Servicebio at Wuhan, China.

### RNA Extraction and qPCR

5 female adult flies of each genotype and αTC1 cells were lysed with Trizol (Thermo Fisher, 15596018) for RNA extraction and cDNA was transcribed using HiScript II Q RT Supermix (Vazyme, R222-01). qPCR was then performed using ChamQ SYBR qPCR Master Mix (Vazyme, Q311-03) on a CFX96 Real-Time System/C1000 Thermal Cycler (Bio-Rad). Expression levels of target genes in fly and mouse were normalized to *RpL32* and *β-actin,* respectively.

The following primers were used in this study.

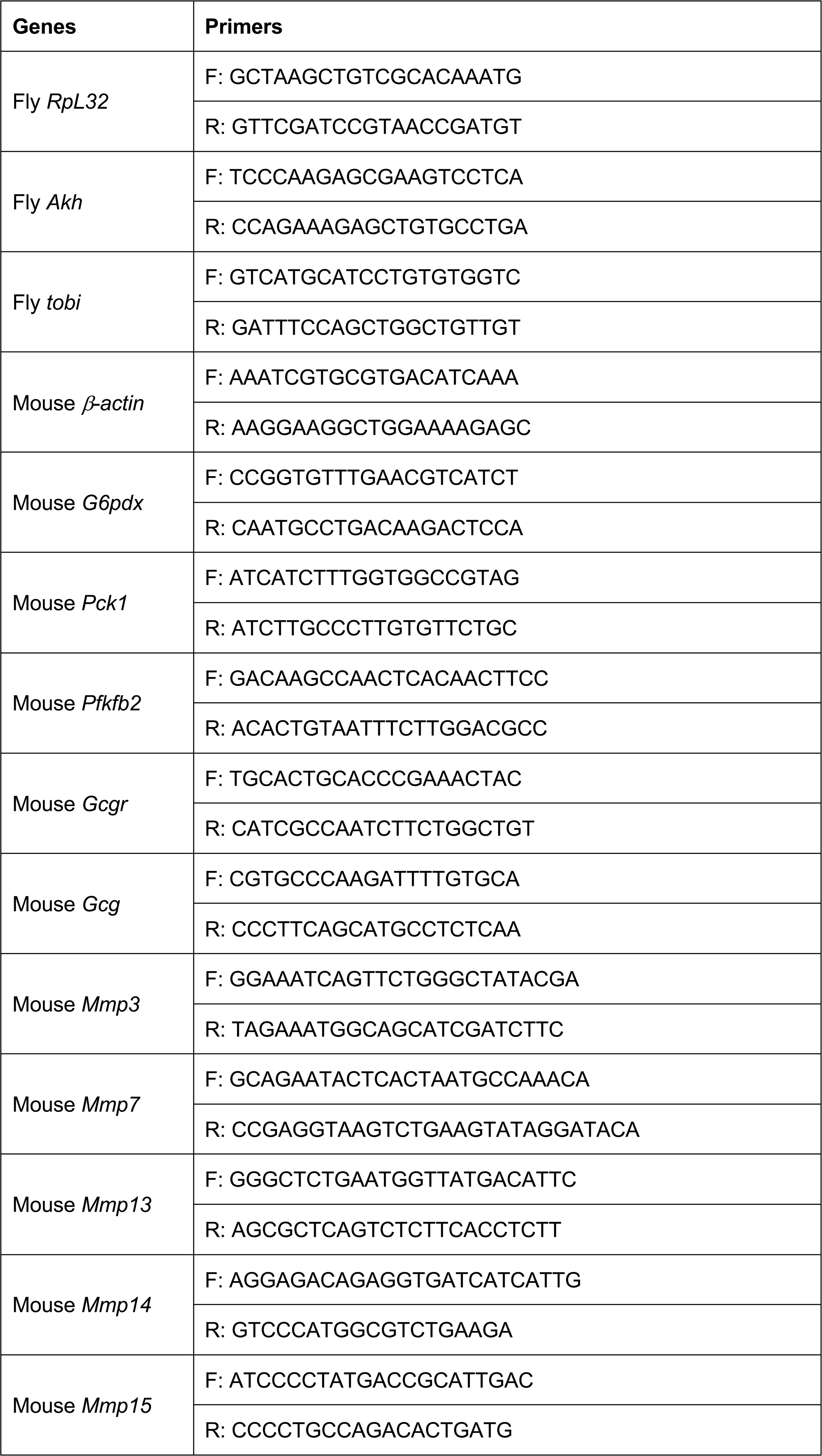
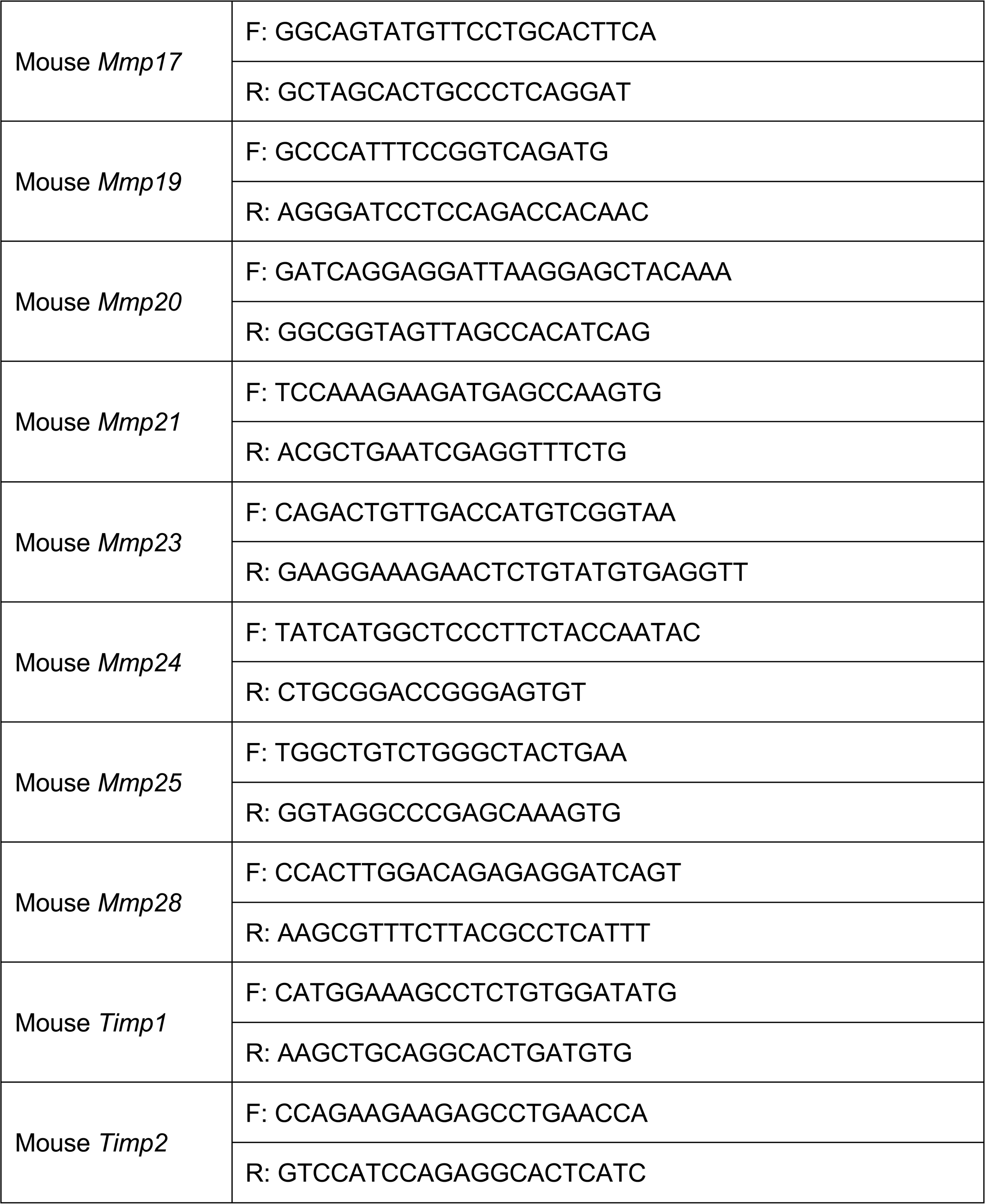

### RNA-seq analysis of gene expression in adult flies

5 female adult flies were dissected for total RNA extraction. After assessing RNA quality with Agilent Bioanalyzer (RIN > 7), mRNAs were enriched by poly-A pull-down. Sequencing libraries were prepared with Illumina Truseq RNA preparation kits and were sequenced using Illumina HiSeq 2000 by Benagen (Wuhan, China). We multiplexed samples in each lane, which yields target number of single-end 100-bp reads for each sample, as a fraction of 180 million reads for the whole lane. After trimming, sequence reads were mapped to the Drosophila genome (FlyBase genome annotation version r6.48) using Tophat. With the uniquely mapped reads, gene expression was quantified using Cufflinks (FPKM values) and HTseq (read counts per gene). Differentially expressed genes were analyzed based on both adjusted *p* value using DSeq2 as well as fold change cut-off.

Prior to fold change calculation, we set to a value of “1” for any FPKM value between 0 and 1 to reduce the possibility that we get large ratio values for genes with negligible levels of detected transcript in both the experimental sample and the wildtype control (e.g. FPKM 0.1 vs. 0.0001), as those ratios are unlikely to have biological relevance. A cut-off of 2-fold change consistently observed among replicates and the adjusted *p* value of 0.1 or lower from DSeq2 analysis were used. Heatmap was generated using MEV_4_7 based on FPKM change.

To analyze the data under biological context, we assembled pathway annotation, including biological processes, KEGG pathways and cellular compartment, from DAVID bioinformatics resources. We calculated enrichment *p* value of gene sets among the up or down regulated genes using hyper-geometric distribution and selected gene sets significantly enrichment with *p* value less than 0.05.

### Measurements of serum glucagon

Serum from blood samples was collected after centrifugation at 3,000 rpm for 15 min at room temperature. 50μL serum was 1:4 diluted with 150 μL dilution buffer to measure glucagon concentration using commercial ELISA kits (Mouse, DGCG0, R&D Systems; human, D711361, Sangon Biotech). All procedures were performed by following the manufacturer’s protocol.

### Glucose Tolerance Test (GTT)

GTTs were performed in 16-hour-fasted male animals as previously reported ^6^. Each animal received an injection of 1 g/Kg glucose in sterile saline. Blood glucose levels were measured at different time points using a glucometer. Mouse forelimb grip strength was measured using Grip Strength Meter (HANDPI, HP-10).

### Statistical Analysis

Data are presented as the mean ± SEM. Unpaired Student’s t test and one-way ANOVA followed by post-hoc test were performed to assess differences. The *P* value of < 0.05 was considered statistically significant.

