## Supplemental Figures for "A tumor-secreted protein utilizes glucagon release to cause host wasting"

### Fig. S1

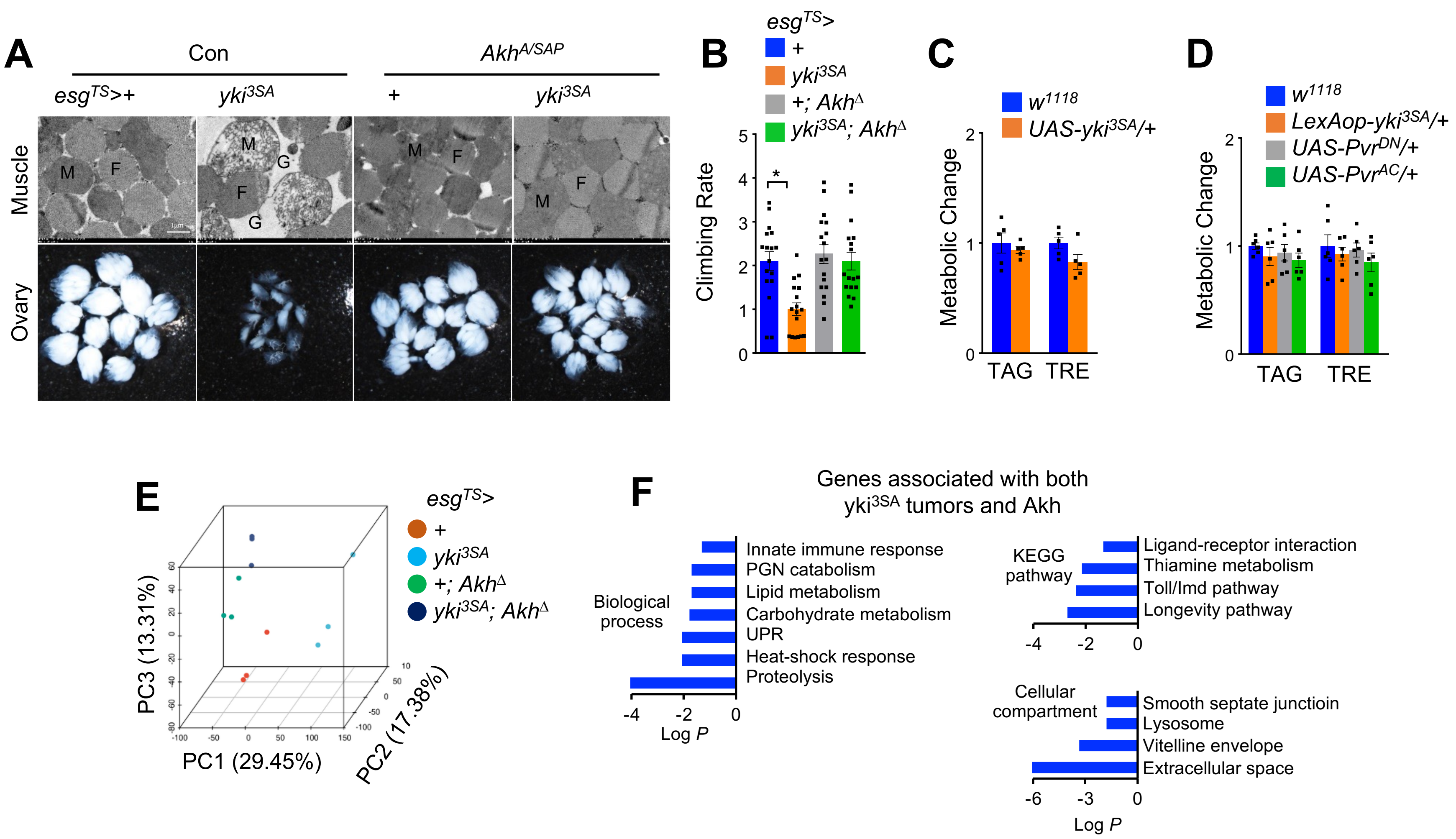

A
Con
$Akh^{A/SAP}$
$esg^{TS}$>+
$yki^{3SA}$
+
$yki^{3SA}$
Muscle
Ovary
M
F
G
B
$esg^{TS}$>
+
$yki^{3SA}$
+; $Akh^{\Delta}$
$yki^{3SA}$; $Akh^{\Delta}$
Climbing Rate
C
$w^{1118}$
UAS-$yki^{3SA}$/+
Metabolic Change
TAG
TRE
D
$w^{1118}$
LexAop-$yki^{3SA}$/+
UAS-$Pvr^{DN}$/+
UAS-$Pvr^{AC}$/+
Metabolic Change
TAG
TRE
E
$esg^{TS}$>
+
$yki^{3SA}$
+; $Akh^{\Delta}$
$yki^{3SA}$; $Akh^{\Delta}$
PC3 (13.31%)
PC1 (29.45%)
PC2 (17.38%)
F
Genes associated with both
$yki^{3SA}$ tumors and Akh
Biological process
Innate immune response
PGN catabolism
Lipid metabolism
Carbohydrate metabolism
UPR
Heat-shock response
Proteolysis
Log P
KEGG pathway
Ligand-receptor interaction
Thiamine metabolism
Toll/Imd pathway
Longevity pathway
Cellular compartment
Smooth septate junctioin
Lysosome
Vitelline envelope
Extracellular space
Log P

Fig. S2

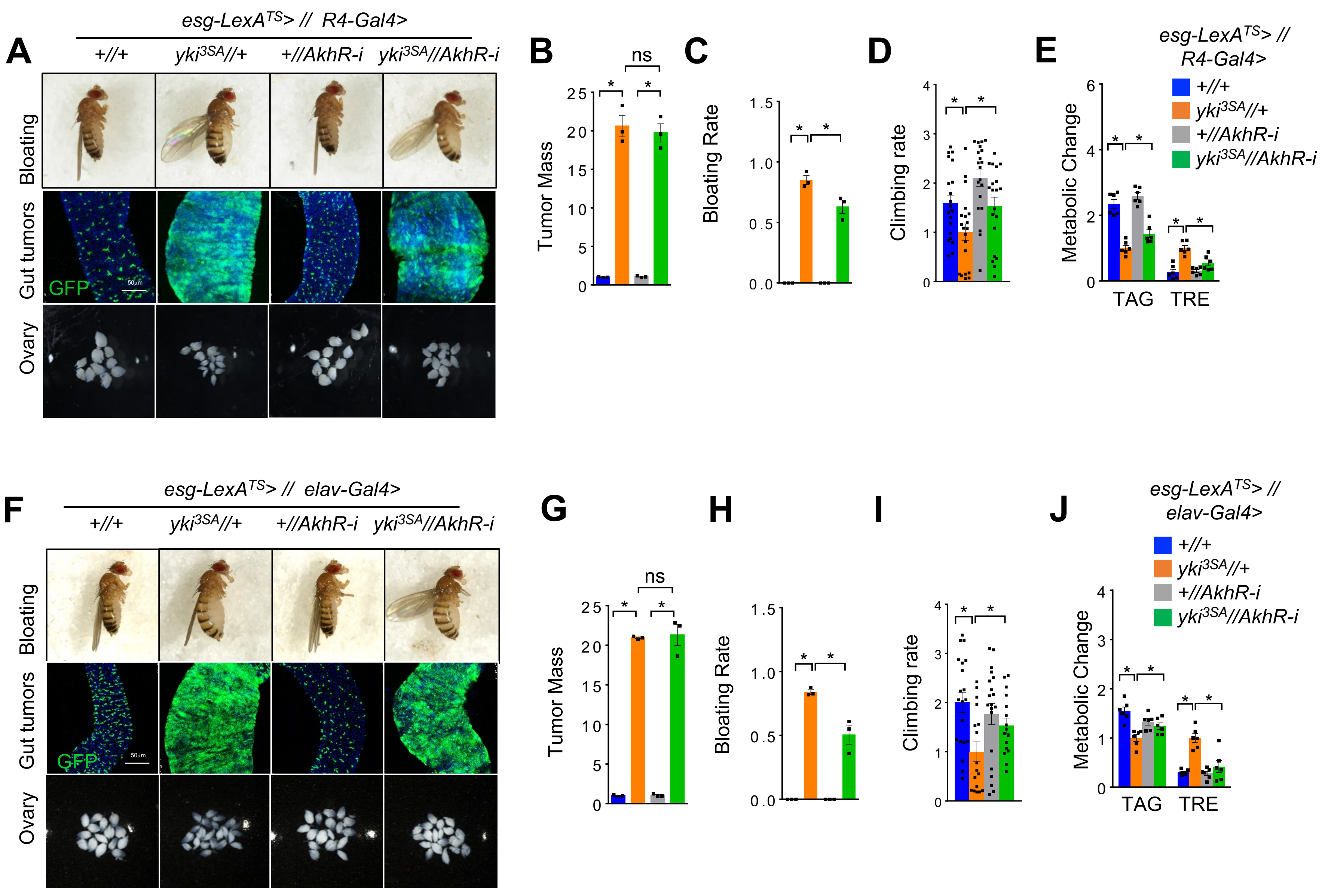
A
esg-LexA$^{TS}$> // R4-Gal4>
+//+
yki$^{3SA}$//+
+//AkhR-i
yki$^{3SA}$//AkhR-i
Bloating
Gut tumors
GFP
50μm
Ovary
B
Tumor Mass
ns
C
Bloating Rate
D
Climbing rate
E
Metabolic Change
TAG
TRE
F
esg-LexA$^{TS}$> // elav-Gal4>
G
H
I
J
esg-LexA$^{TS}$> // elav-Gal4>

### Fig. S3

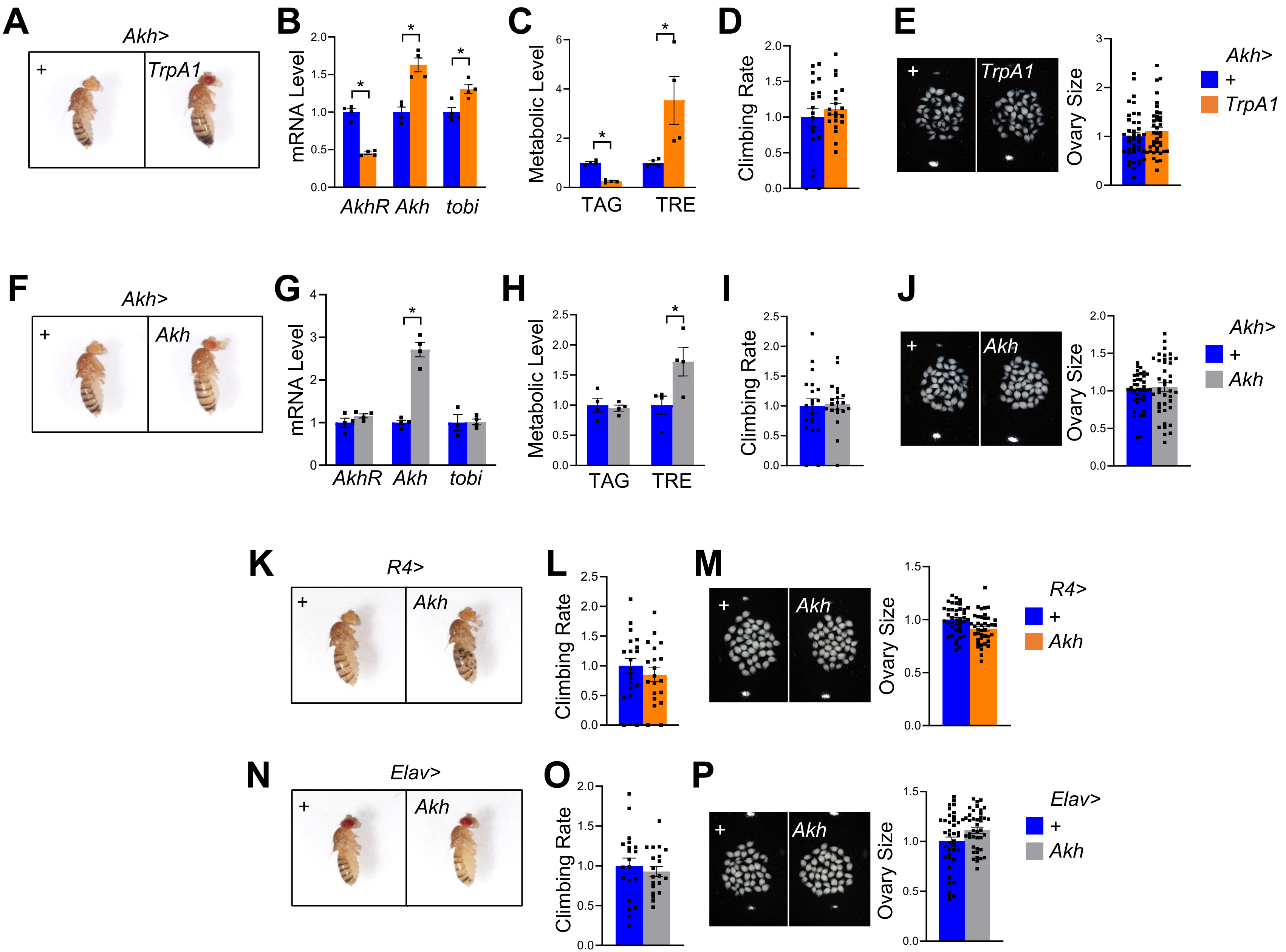

A
Akh>
+
TrpA1
B
mRNA Level
AkhR Akh tobi
C
Metabolic Level
TAG
TRE
D
Climbing Rate
E
Ovary Size
Akh>
+
TrpA1
F
Akh>
+
Akh
G
mRNA Level
AkhR Akh tobi
H
Metabolic Level
TAG
TRE
I
Climbing Rate
J
Ovary Size
Akh>
+
Akh
K
R4>
+
Akh
L
Climbing Rate
M
Ovary Size
R4>
+
Akh
N
Elav>
+
Akh
O
Climbing Rate
P
Ovary Size
Elav>
+
Akh

**Fig. S4**

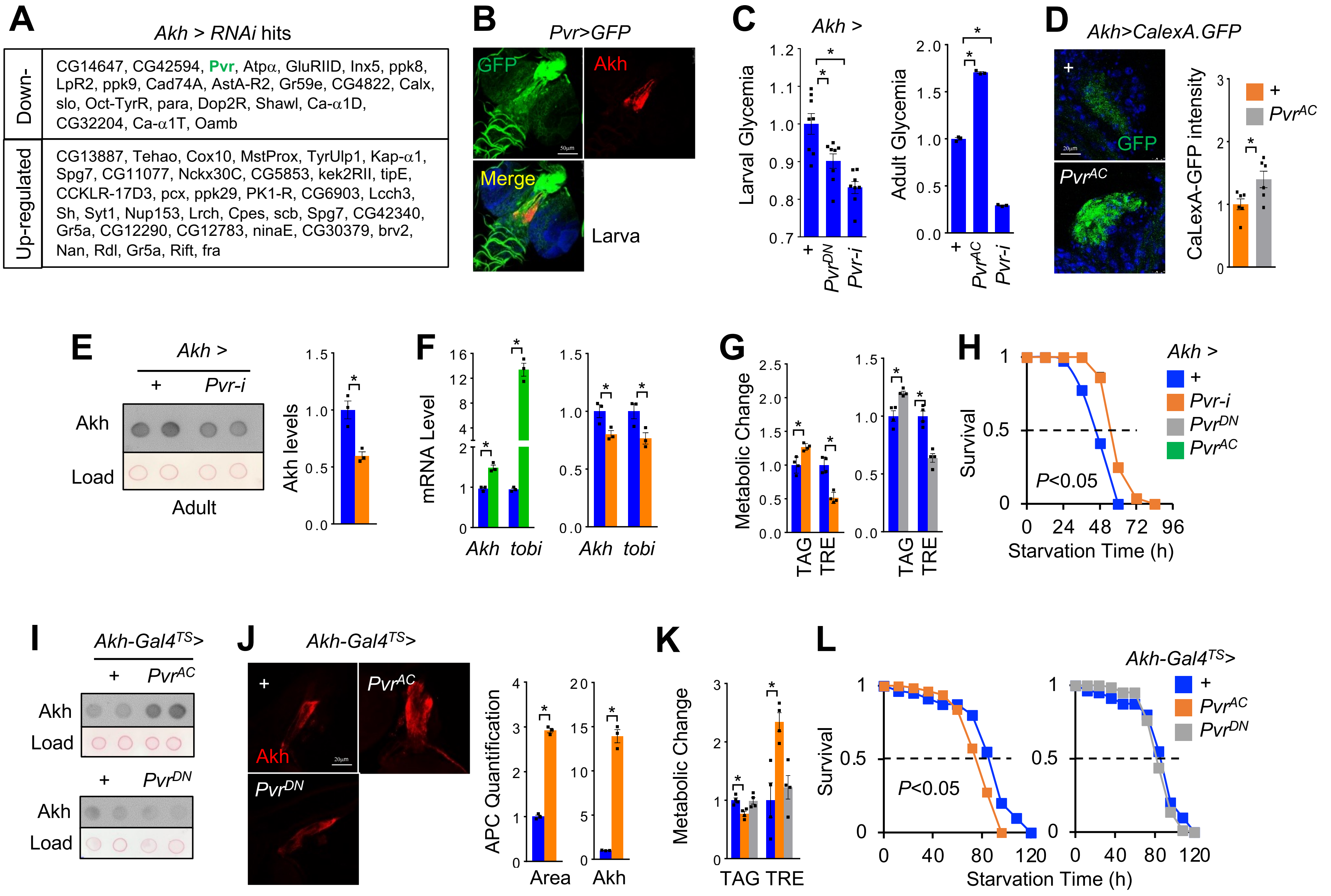
A
Akh > RNAi hits
Down-
CG14647, CG42594, Pvr, Atpα, GluRIID, Inx5, ppk8, LpR2, ppk9, Cad74A, AstA-R2, Gr59e, CG4822, Calx, slo, Oct-TyrR, para, Dop2R, Shawl, Ca-α1D, CG32204, Ca-α1T, Oamb
Up-regulated
CG13887, Tehao, Cox10, MstProx, TyrUlp1, Kap-α1, Spg7, CG11077, Nckx30C, CG5853, kek2RII, tipE, CCKLR-17D3, pcx, ppk29, PK1-R, CG6903, Lcch3, Sh, Syt1, Nup153, Lrch, Cpes, scb, Spg7, CG42340, Gr5a, CG12290, CG12783, ninaE, CG30379, brv2, Nan, Rdl, Gr5a, Rift, fra
B
Pvr>GFP
GFP
Akh
Merge
Larva
C
Akh >
Larval Glycemia
Adult Glycemia
+
PvrDN
Pvr-i
PvrAC
D
Akh>CalexA.GFP
GFP
PvrAC
CaLexA-GFP intensity
E
Akh >
Pvr-i
Akh
Load
Adult
Akh levels
F
mRNA Level
Akh
tobi
G
Metabolic Change
TAG
TRE
H
Survival
P<0.05
Starvation Time (h)
I
Akh-Gal4TS>
J
Akh-Gal4TS>
APC Quantification
Area
K
L
Akh-Gal4TS>
Survival
P<0.05
Starvation Time (h)

### Fig. S5

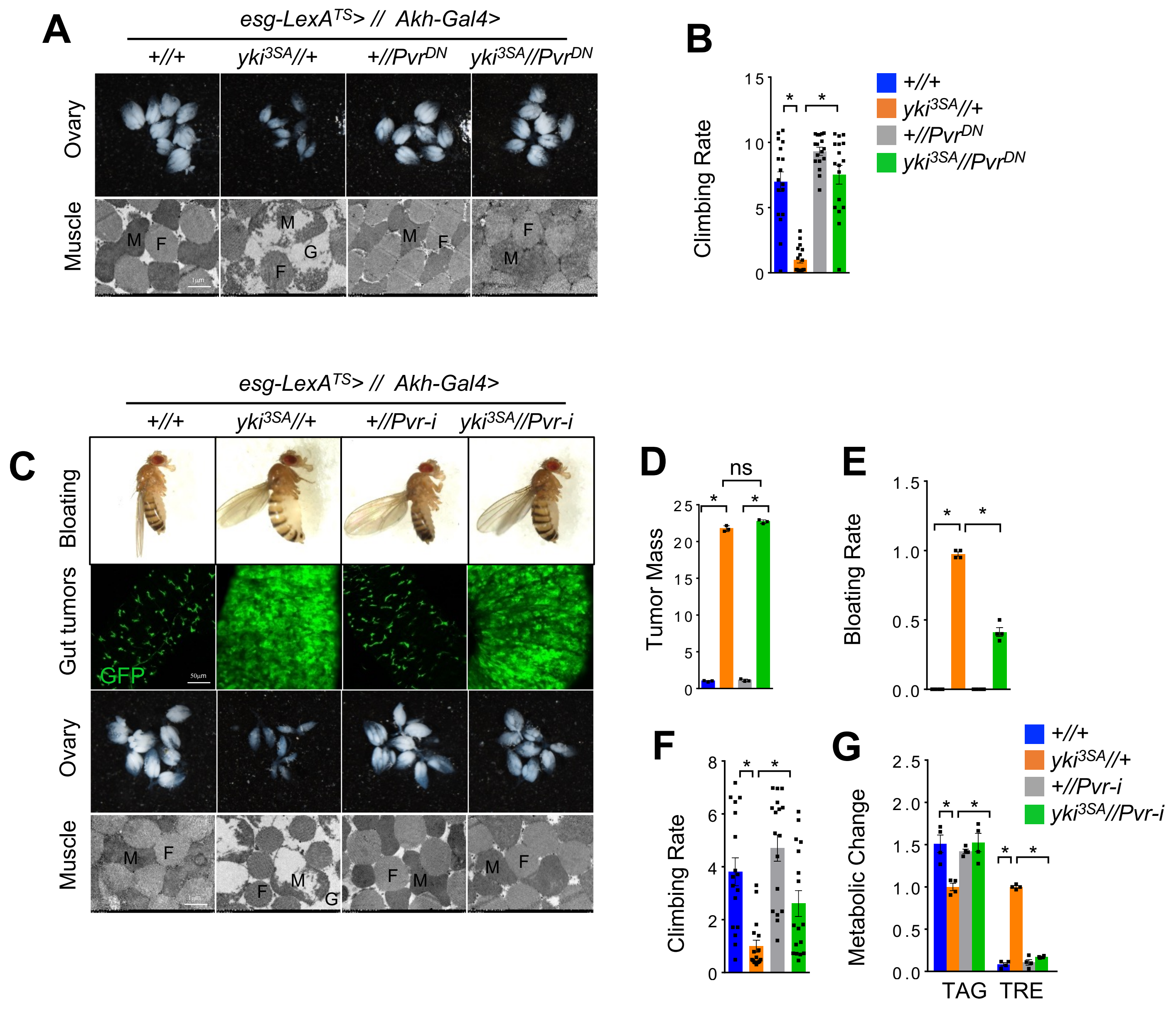
A
esg-LexA^TS> // Akh-Gal4>
+//+
yki^3SA//+
+//Pvr^DN
yki^3SA//Pvr^DN
Ovary
Muscle
M
F
G
1μm
B
Climbing Rate
15
10
5
0
*
+//+
yki^3SA//+
+//Pvr^DN
yki^3SA//Pvr^DN
esg-LexA^TS> // Akh-Gal4>
+//+
yki^3SA//+
+//Pvr-i
yki^3SA//Pvr-i
C
Bloating
Gut tumors
GFP
50μm
Ovary
Muscle
D
Tumor Mass
ns
25
20
15
10
5
0
E
Bloating Rate
1.5
1.0
0.5
0.0
F
Climbing Rate
8
6
4
2
0
G
Metabolic Change
2.5
2.0
1.5
1.0
0.5
0.0
TAG
TRE
+//+
yki^3SA//+
+//Pvr-i
yki^3SA//Pvr-i

**Fig. S6**

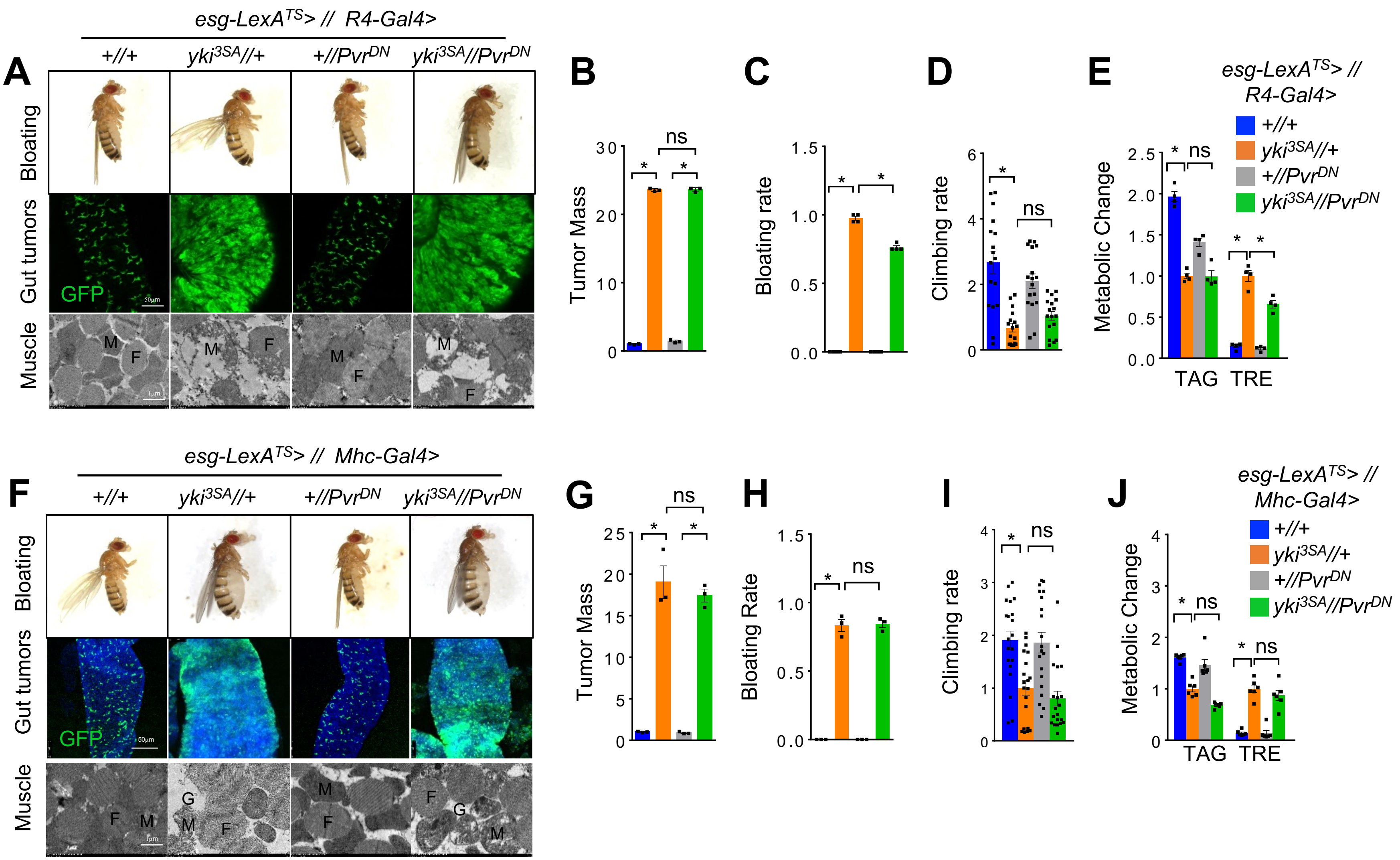
esg-LexA^TS> // R4-Gal4>
A
+//+
yki3SA//+
+//PvrDN
yki3SA//PvrDN
Bloating
Gut tumors
GFP
50µm
Muscle
M
F
1µm
B
Tumor Mass
30
20
10
0
ns
*
C
Bloating rate
1.5
1.0
0.5
0.0
D
Climbing rate
6
4
2
0
E
Metabolic Change
2.5
2.0
1.5
1.0
0.5
0.0
TAG
TRE
esg-LexA^TS> // R4-Gal4>
esg-LexA^TS> // Mhc-Gal4>
F
G
H
Bloating Rate
I
J
25
15
5
3
1
G
esg-LexA^TS> // Mhc-Gal4>

Fig. S7

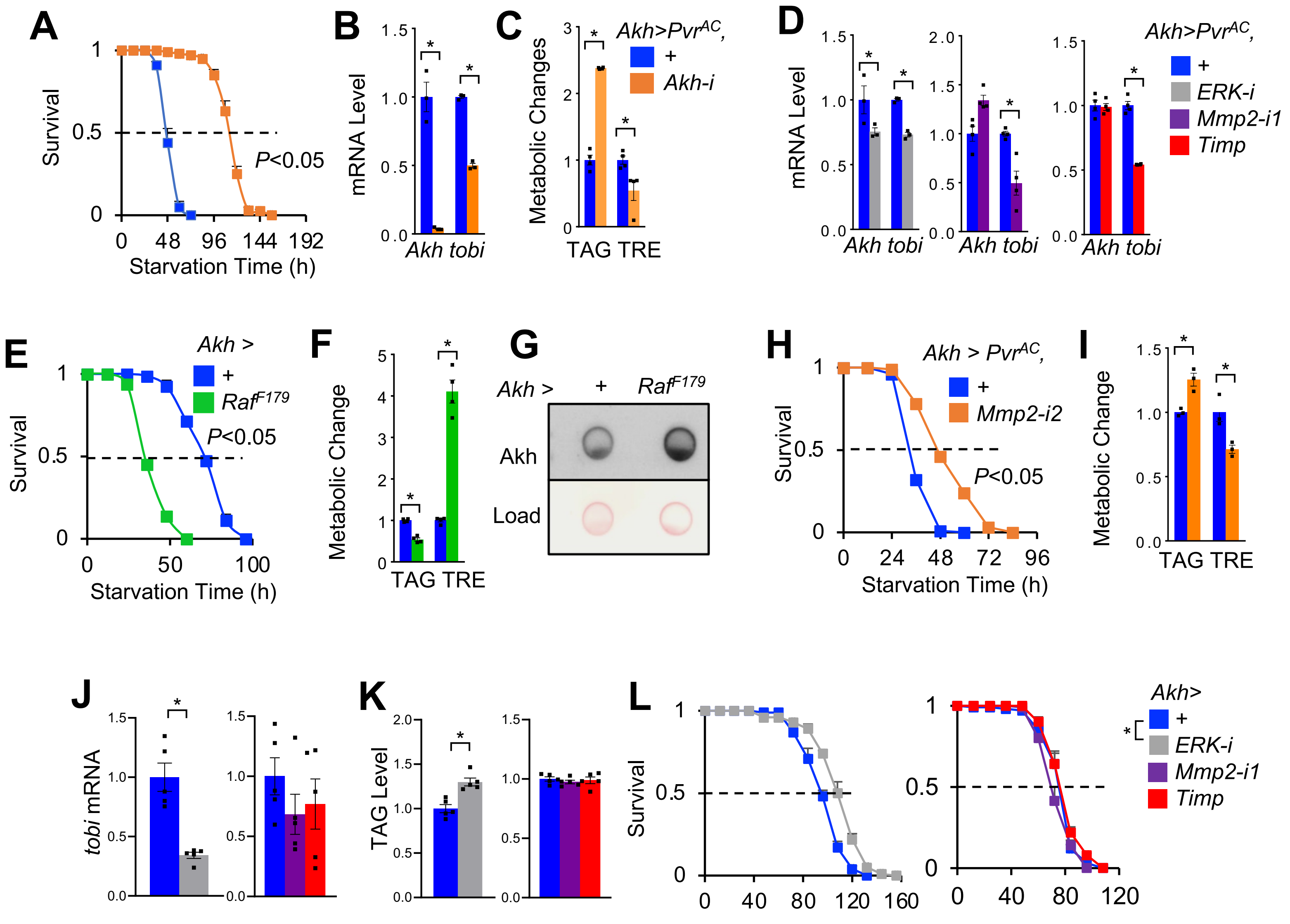

A
Survival
Starvation Time (h)
P<0.05
B
mRNA Level
Akh tobi
C
Metabolic Changes
TAG TRE
Akh>PvrAC,
+
Akh-i
D
mRNA Level
Akh tobi
Akh>PvrAC,
+
ERK-i
Mmp2-i1
Timp
E
Survival
Starvation Time (h)
Akh >
+
RafF179
P<0.05
F
Metabolic Change
TAG TRE
G
Akh >
+
RafF179
Akh
Load
H
Survival
Starvation Time (h)
Akh > PvrAC,
+
Mmp2-i2
P<0.05
I
Metabolic Change
TAG TRE
J
tobi mRNA
K
TAG Level
L
Survival
Akh>
+
ERK-i
Mmp2-i1
Timp

**Fig. S8**

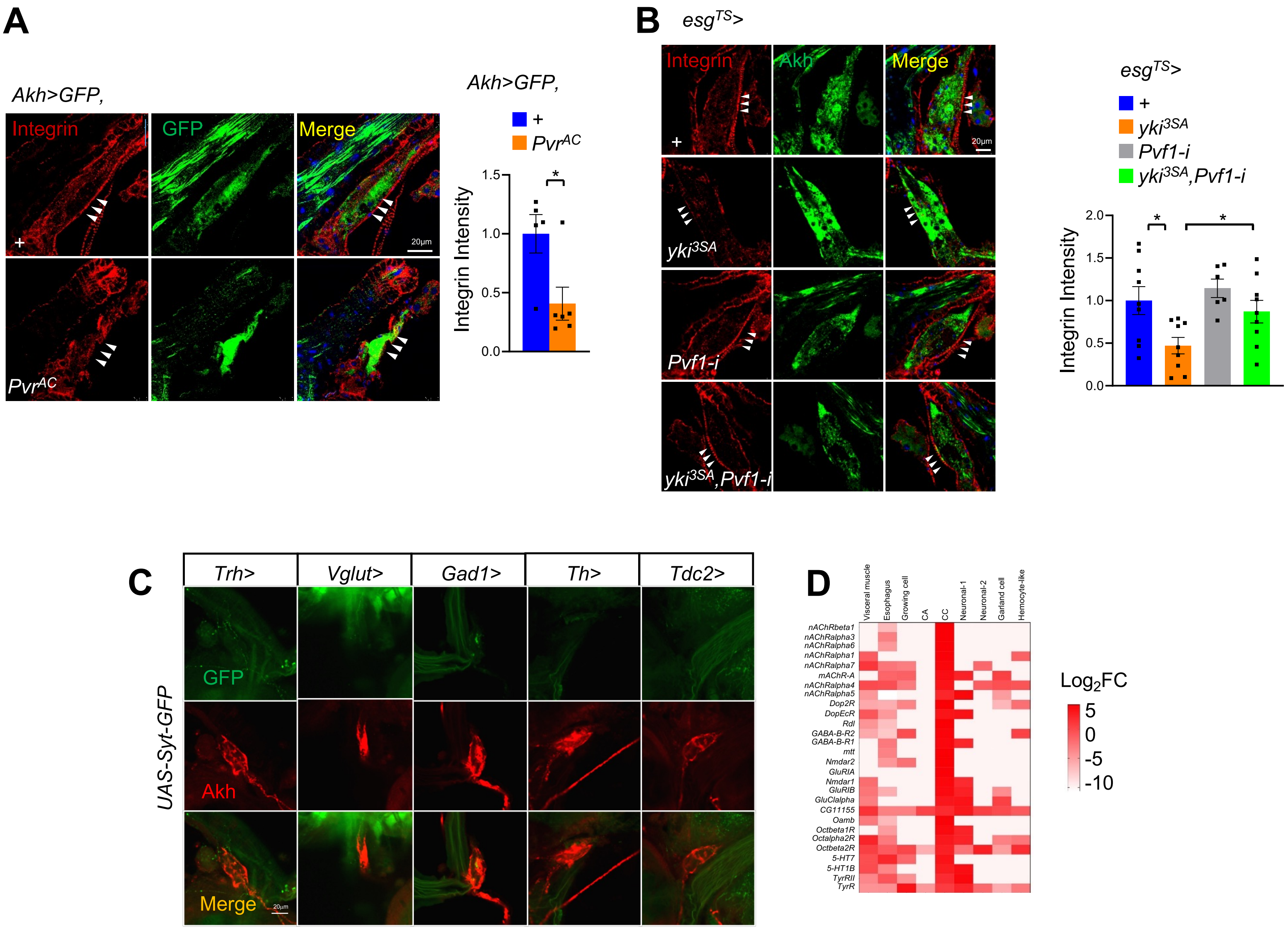
A
Akh>GFP,
Integrin
GFP
Merge
+
PvrAC
20µm
Akh>GFP,
+
PvrAC
Integrin Intensity
1.5
1.0
0.5
0.0
*
B
esgTS>
Integrin
Akh
Merge
+
yki3SA
Pvf1-i
yki3SA,Pvf1-i
20µm
esgTS>
+
yki3SA
Pvf1-i
yki3SA,Pvf1-i
Integrin Intensity
2.0
1.5
1.0
0.5
0.0
*
*
C
Trh>
Vglut>
Gad1>
Th>
Tdc2>
UAS-Syt-GFP
GFP
Akh
Merge
20µm
D
Visceral muscle
Esophagus
Growing cell
CA
CC
Neuronal-1
Neuronal-2
Garland cell
Hemocyte-like
nAChRbeta1
nAChRalpha3
nAChRalpha6
nAChRalpha1
nAChRalpha7
mAChR-A
nAChRalpha4
nAChRalpha5
Dop2R
DopEcR
Rdl
GABA-B-R2
GABA-B-R1
mtt
Nmdar2
GluRIA
Nmdar1
GluRIB
GluClalpha
CG11155
Oamb
Octbeta1R
Octalpha2R
Octbeta2R
5-HT7
5-HT1B
TyrRII
TyrR
Log2FC
5
0
-5
-10

Fig. S9

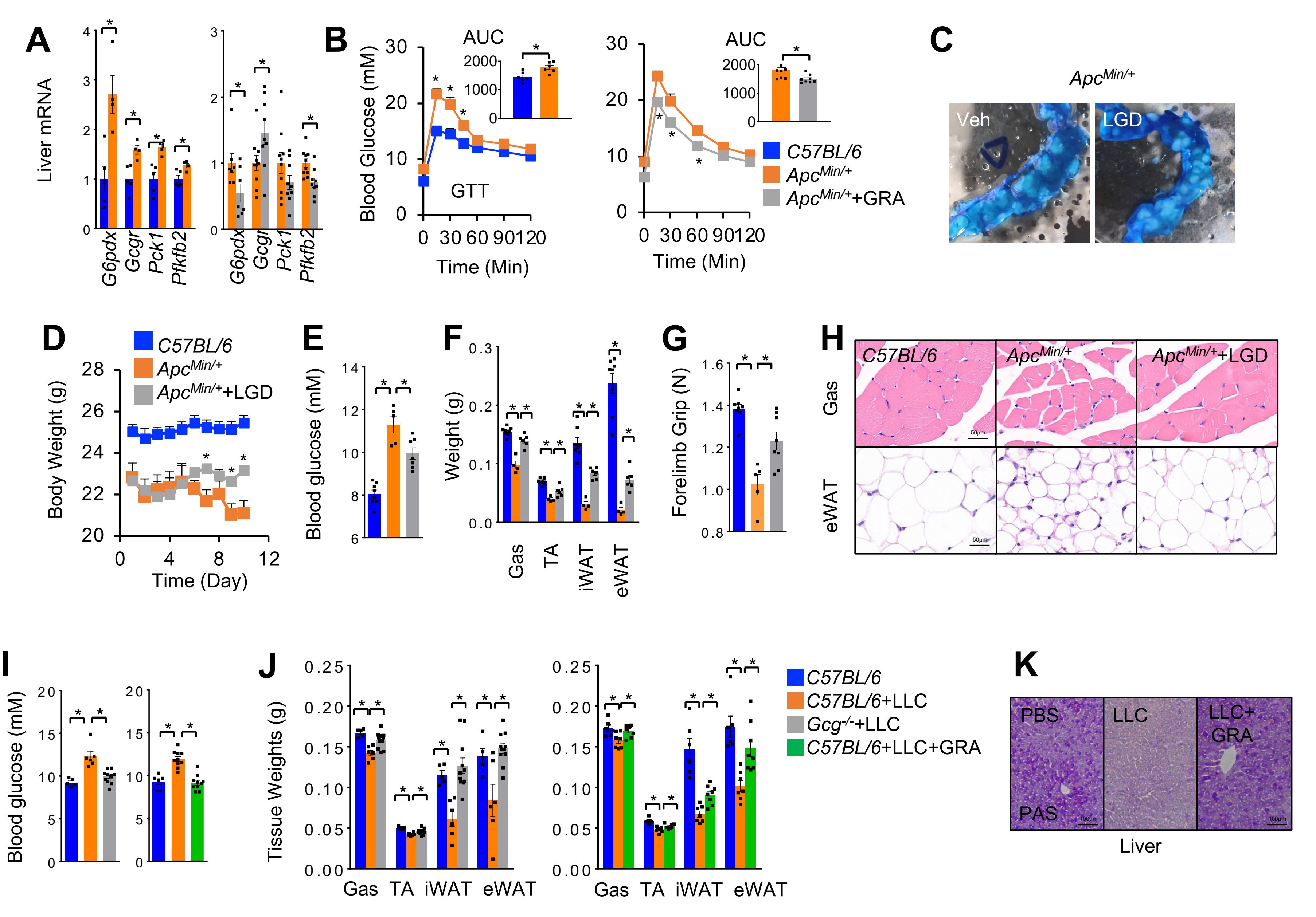
A
Liver mRNA
G6pdx
Gcgr
Pck1
Pfkfb2
B
Blood Glucose (mM)
AUC
GTT
Time (Min)
C57BL/6
ApcMin/+
ApcMin/++GRA
C
ApcMin/+
Veh
LGD
D
Body Weight (g)
Time (Day)
C57BL/6
ApcMin/+
ApcMin/++LGD
E
Blood glucose (mM)
F
Weight (g)
Gas
TA
iWAT
eWAT
G
Forelimb Grip (N)
H
C57BL/6
ApcMin/+
ApcMin/++LGD
Gas
eWAT
I
Blood glucose (mM)
J
Tissue Weights (g)
Gas
TA
iWAT
eWAT
C57BL/6
C57BL/6+LLC
Gcg-/-+LLC
C57BL/6+LLC+GRA
K
PBS
LLC
LLC+ GRA
PAS
Liver

### Fig. S10

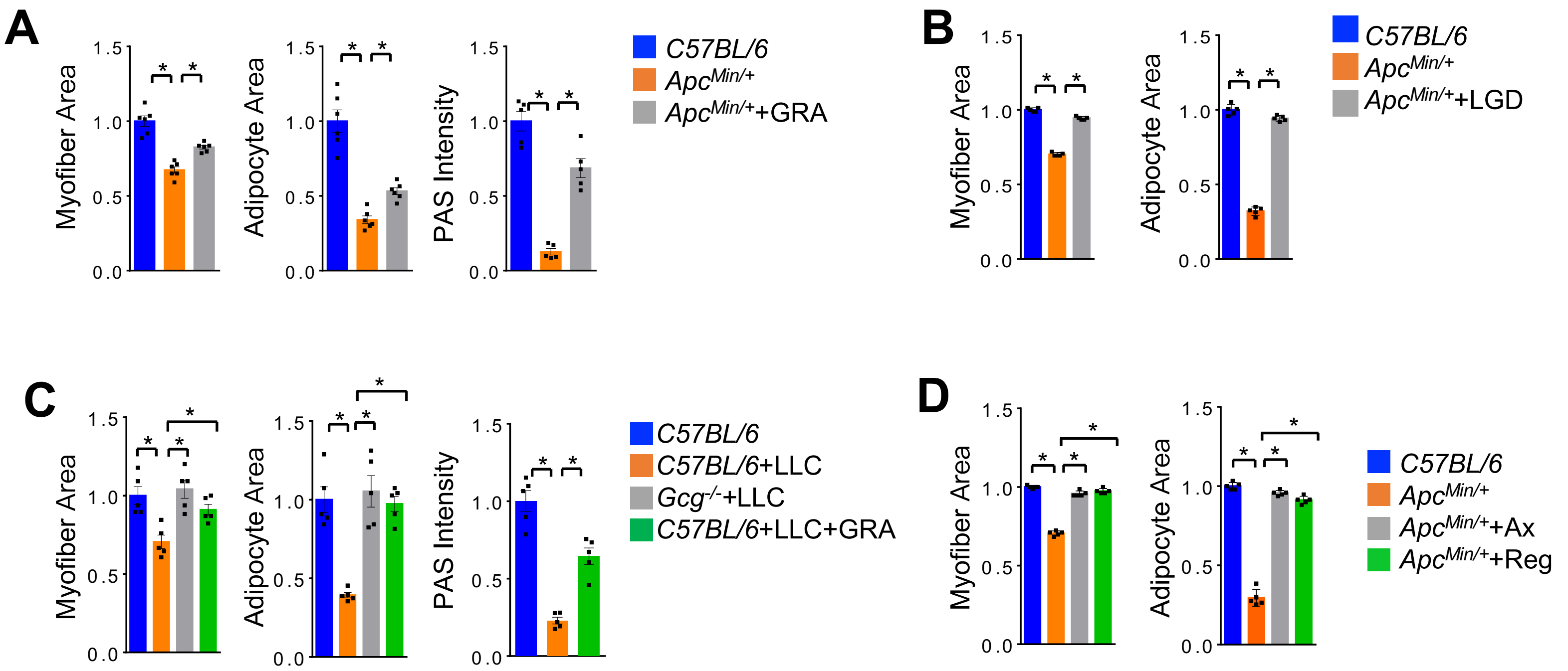


**E** GSE54973 (RPKM)

| Gene | α1 | α2 | β1 | β2 |
|---|---|---|---|---|
| *Flt1/Vegfr1* | 0.95 | 0.80 | 1.36 | 1.89 |
| *Kdr/Vegfr2* | 3.69 | 3.33 | 5.95 | 6.06 |
| *Flt4/Vegfr3* | 0.62 | 0.68 | 0.72 | 0.69 |
| *Pdgfra* | 1.99 | 3.09 | 0.02 | 0.01 |
| *Pdgfrb* | 3.25 | 4.34 | 0.83 | 0.95 |
| *Chrm1/AchR-M1* | 0.06 | 0.16 | 0.01 | 0.00 |
| *Chrm2/AchR-M2* | 0.02 | 0.00 | 0.00 | 0.00 |
| *Chrm3/AchR-M3* | 1.95 | 4.43 | 14.49 | 13.65 |
| *Chrm4/AchR-M4* | 6.53 | 6.63 | 2.53 | 2.89 |
| *Chrm5/AchR-M5* | 0.00 | 0.00 | 0.00 | 0.00 |

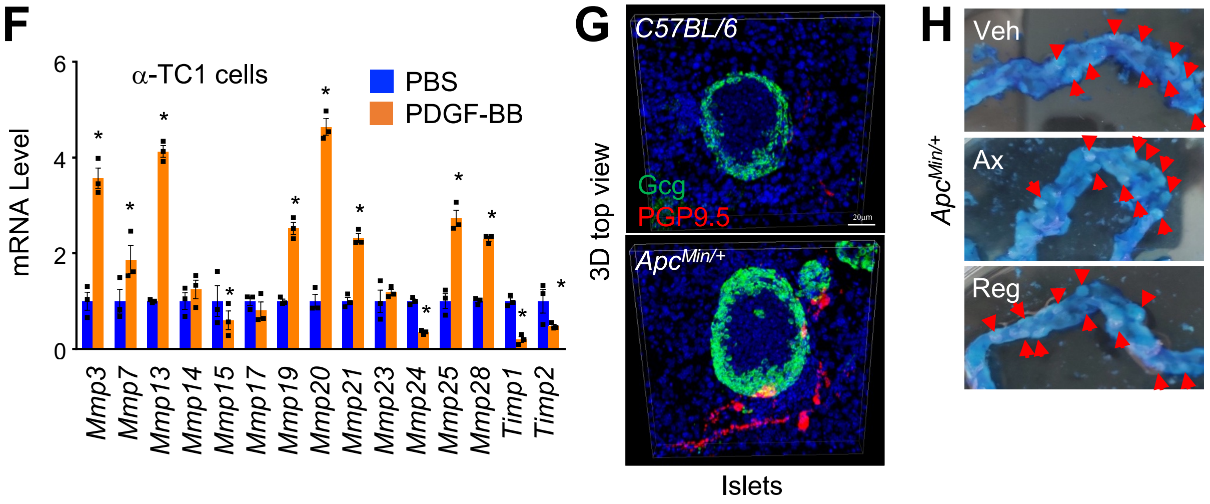
